# Predicting SARS-CoV-2 evolution dynamics with spatiotemporal resolution by DMS-empowered protein language model

**DOI:** 10.1101/2025.10.17.683094

**Authors:** Sijie Yang, Xiaowei Luo, Jiejian Luo, Fanchong Jian, Yunlong Cao

## Abstract

Early identification of emerging dominant SARS-CoV-2 variants is essential for effective pandemic preparedness, yet existing methodologies face significant limitations. Experimental characterizations are costly and not feasible for real-time surveillance, whereas existing computational approaches cannot achieve satisfactory precision in predicting future dominant lineages and fail to capture the spatiotemporal dynamics of fitness under evolving host immune pressures. Here, we introduce DeepCoV (DMS-Empowered Evolution Prediction of CoronaVirus), a deep-learning framework for the dynamic identification of novel variants with high potential to become prevelent. It integrates deep mutational scanning (DMS)-derived mutation phenotypes with epidemiological surveillence data reflecting historical viral evolution and the dynamic fitness landscape. DeepCoV accurately forecasted the dominance of recently circulating lineages a month in advance, achieving a 90% reduction in false discovery rate while capturing temporal and geographic dynamics of variant spread and reconstructing their regional prevalence trajectories. Moreover, DeepCoV identified mutational hotspots of Omicron-derived backbones *in silico*, revealing convergent evolution trends. This scalable solution enables timely identification of immune-evasive variants and prospective alert of critical mutations, providing actionable insights for vaccine updates and pandemic surveillance.

## Main

The evolutionary arms race between pathogens and human immunity emphasizes the necessity of proactive surveillance of emerging variants^1–3^. For rapidly evolving viruses such as SARS-CoV-2, early identification of high-growth lineages is essential to pandemic resilience, enabling timely updates to vaccines and informing the development of antibody-based therapeutics^4–45^.

Although high-throughput experimental approaches such as DMS can generate valuable data and insights into the functional impact of individual mutations, their substantial resource requirements restrict their application for continuous surveillance^46–52^. Moreover, DMS-based methods are inherently incapable of capturing the evolutionary dynamics of full viral sequences within populations, as they typically probe only a subdomain of the full spike protein or restricted set of mutations and face challenges in modeling epistatic interactions, given the prohibitively large mutational combinatorial space^51,53^. These methods proved critical during the early phases of the COVID-19 pandemic but have become increasingly impractical.

Statistical models based on sequence frequency dynamics—such as linear growth advantage estimation—offer alternative tools for inferring variant fitness directly based on epidemiological surveillance^54^. However, their predictive reliability declines substantially when data remain sparse, especially during the early stage of novel lineage emergence and in the post-pandemic period, when sequencing efforts have markedly decreased. More sophisticated frameworks, including EpiScore and PyR0, incorporate evolutionary constraints by modeling sequence prevalence over time^55,56^. However, they often struggle to pinpoint the most prevalent circulating strains. Without incorporating sequence information, such epidemiological analyses remain largely phenomenological and offer limited mechanistic insight into why certain variants rise to dominance while others fade. Meanwhile, reliably capturing sequence-level features remains inherently challenging for statistical approaches.

Artificial intelligence (AI)-based methods have thus emerged as promising tools for forecasting viral evolution^56–65^. AI-based approaches can overcome the combinatorial explosion arising from multiple mutations within viral sequences and enable the integrated learning of large, diverse strain sets, including capturing amino acid sequence evolutionary patterns. However, existing models remain limited in their ability to prospectively identify emerging dominant variants with sufficient accuracy. Viral protein sequence or structure-based approaches such as EVEscape (variational autoencoder based) and TEMPO (Transformer-based) exhibit strong representational capacity but typically overlook functional data, particularly experimentally derived measurements of antibody escape and other virological phenotypes captured through DMS^58,65^. Methods such as E2VD and CoVFit have advanced by leveraging mutational phenotypes, but they still often neglect the dynamic host immune context, which is critical for capturing the spatiotemporal dimensions of viral transmission^62,66^. Moreover, most current computational models fail to capture the dynamic viral fitness landscape under the co-evolution of host immune pressures, frequently underperforming compared to even simple linear growth advantage estimators in real-world predictive applications.

Despite recent progress in both experimental and computational approaches, a major challenge remains in jointly capturing the spatiotemporal dynamic fitness landscape for viral evolution under evolving population immune pressures or herd immunity. Even within the same lineage, transmission advantages can diverge substantially across regions and time with distinct immune history. For instance, the Omicron sublineage XBB.1.5 exhibited markedly different growth trajectories in North America and parts of Asia, where preexisting immunity was shaped predominantly by prior BA.5 or BA.2.75 infections, respectively, highlighting how regional immune histories can modulate the apparent fitness of otherwise genetically similar variants. Meanwhile, DMS has been underutilized for predictive purposes, as most applications have remained descriptive, focusing on characterizing escape mutations rather than integrating functional data into dynamic evolutionary modeling^46–49,51^.

To bridge this gap, building on our long-term experience and systematic understanding of DMS on viral antigens, we developed DeepCoV (DMS-Empowered Evolution Prediction of CoronaVirus), a predictive framework that integrates DMS-derived functional phenotypes, evolutionary sequence information, and epidemiological data reflecting immune pressures in human populations. By leveraging Transformer-based architectures, DeepCoV learns the mechanistic relationships between mutation effects and variant fitness, while incorporating background epidemiological data and related sequence with pretrained protein language models to accurately model viral evolution at spatiotemporal resolution^3,64,67^. Collectively, DeepCoV provides a scalable and biologically grounded framework for forecasting SARS-CoV-2 evolutionary trajectories, thereby enhancing global preparedness and informing timely public health interventions.

## Results

### DeepCoV architecture

Accurately forecasting the evolutionary dynamics of SARS-CoV-2 requires integrating information that reflects both the intrinsic viral fitness for infection and transmission, and the impacts of population immune pressure. To this end, we designed DeepCoV, a neural network framework that predicts the future prevelence of any SARS-CoV-2 Spike or RBD variant, leveraging three complementary data collected during months before the time of prediction: 1) Mutiple sequence alignment (MSA) of viral antigen sequences including the variant for prediction and other co-circulating strains that prevail and compete within the same environment. 2) The proportions of the above strains since 180 days before the day of prediction, capturing the recent evolution of viral fitness with the population-level immunity and selective pressure considered implicitly, and endowing the model with the ability to learn the spatiotemporal dynamics of variant circulation. 3) Auxiliary functional mutation phenotypes derived from DMS quantify the impacts on virological characteristics and antigenicity of single mutations on the antigen, thereby grounding the model in experimentally validated datasets and molecular mechanisms. (Fig. 1 and Extended Data Fig. 1). For sequence modeling, we implemented an evolutionary module that captures amino acid substitution patterns in the target sequence and the most prevalent sequences at the time using the pretrained ESM-MSA-1b model, which learns evolutionary constraints from MSA^68^. To incorporate temporal dynamics, the historical proportion embedder employs a long short-term memory (LSTM) network to model prevalence trajectories over a sliding window spanning recent months (typically 180 days) ^69^. These sequence and prevalence representations are concatenated and passed through an axial-attention module to capture residue-prevalence dependencies^64^. This architecture inherently encodes population-level immune histories by integrating background sequence context and prevalence dynamics, thereby reflecting how prior infections and vaccinations shape the viral fitness landscape and influence variant emergence. Finally, the DMS processor incorporates quantitative mutational phenotypes, including antibody escape, human antisera evasion, ACE2 binding affinity, and protein expression, offering mechanistic insights into variant viability and transmissibility^4,34,46–51,70–77^. By integrating these heterogeneous data streams, DeepCoV learns how individual amino acid substitutions and their functional consequences translate into shifts in population-level prevalence, thereby linking molecular evolution with epidemiological outcomes. This unified representation enables the model to infer variant fitness, anticipate lineage competition outcomes, and forecast regional prevalence trends over future time windows. Finally, DeepCoV is expected to predict the future proportion of a certain strain at any time point, with the sequences of itself and other co-circulating strains, their historical proportions, and the impacts of mutations carried by the strain from DMS as inputs.

**Figure 1.**
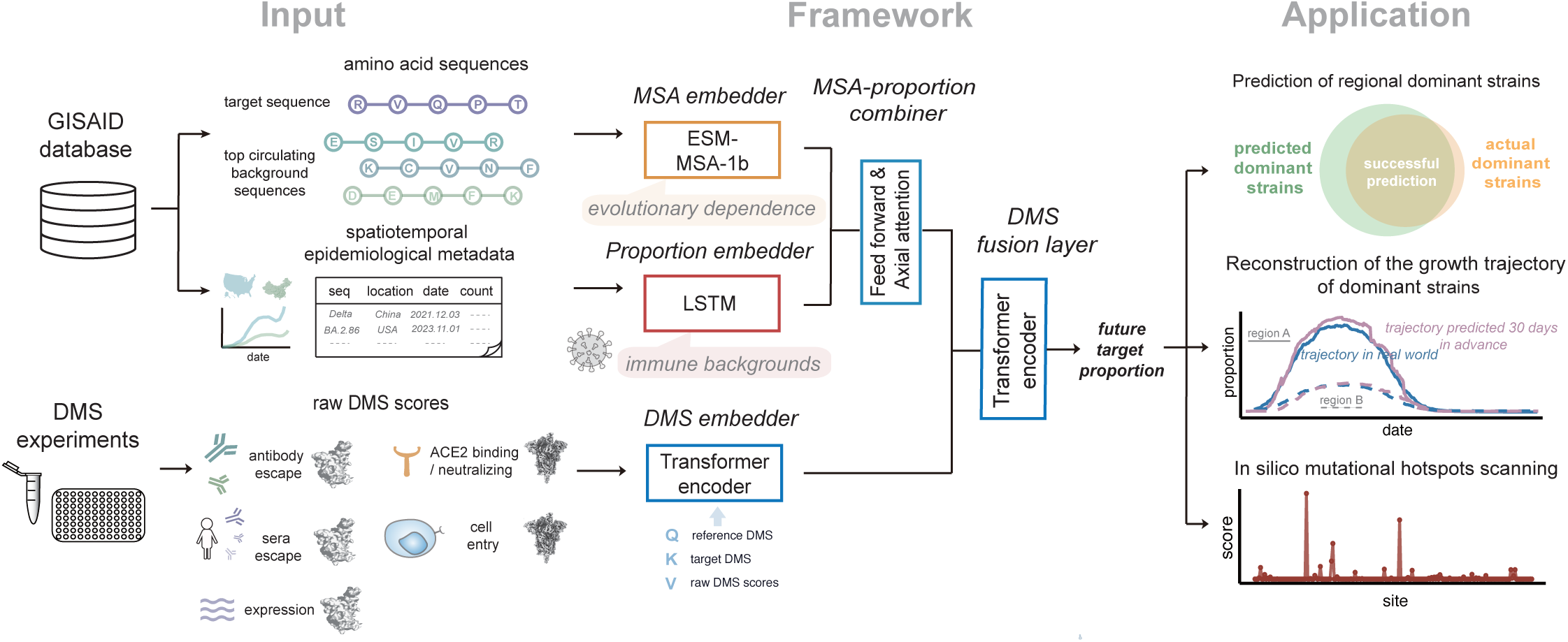
Overview of model architecture and predictive applications. The framework integrates sequence evolution, epidemiological information, and DMS data for variant dominance prediction, reconstruction of growth trajectory and mutational hotspots scanning. The emergence date of JN.1 is defined as the earliest date on which the number of JN.1 RBD sequences exceeds 10.

### DeepCoV accurately predicts predominant variants

We first evaluated DeepCoV for the early identification of emerging dominant JN.1 variants using a retrospective approach^17,19,38,78–83^. Specifically, the model was trained exclusively on epidemiological records and receptor binding domain (RBD) sequences collected prior to October 2023, along with DMS profiles generated before the emergence of JN.1 (Fig. 2a). To prioritize learning from dominant lineages, low-prevalence variants were filtered out from the training dataset. The remaining RBD sequences were then randomly assigned to the training and validation sets at a 9:1 ratio, ensuring that all members of a given cluster were confined to the same split and preventing data leakage due to temporal dependencies.

**Figure 2.**
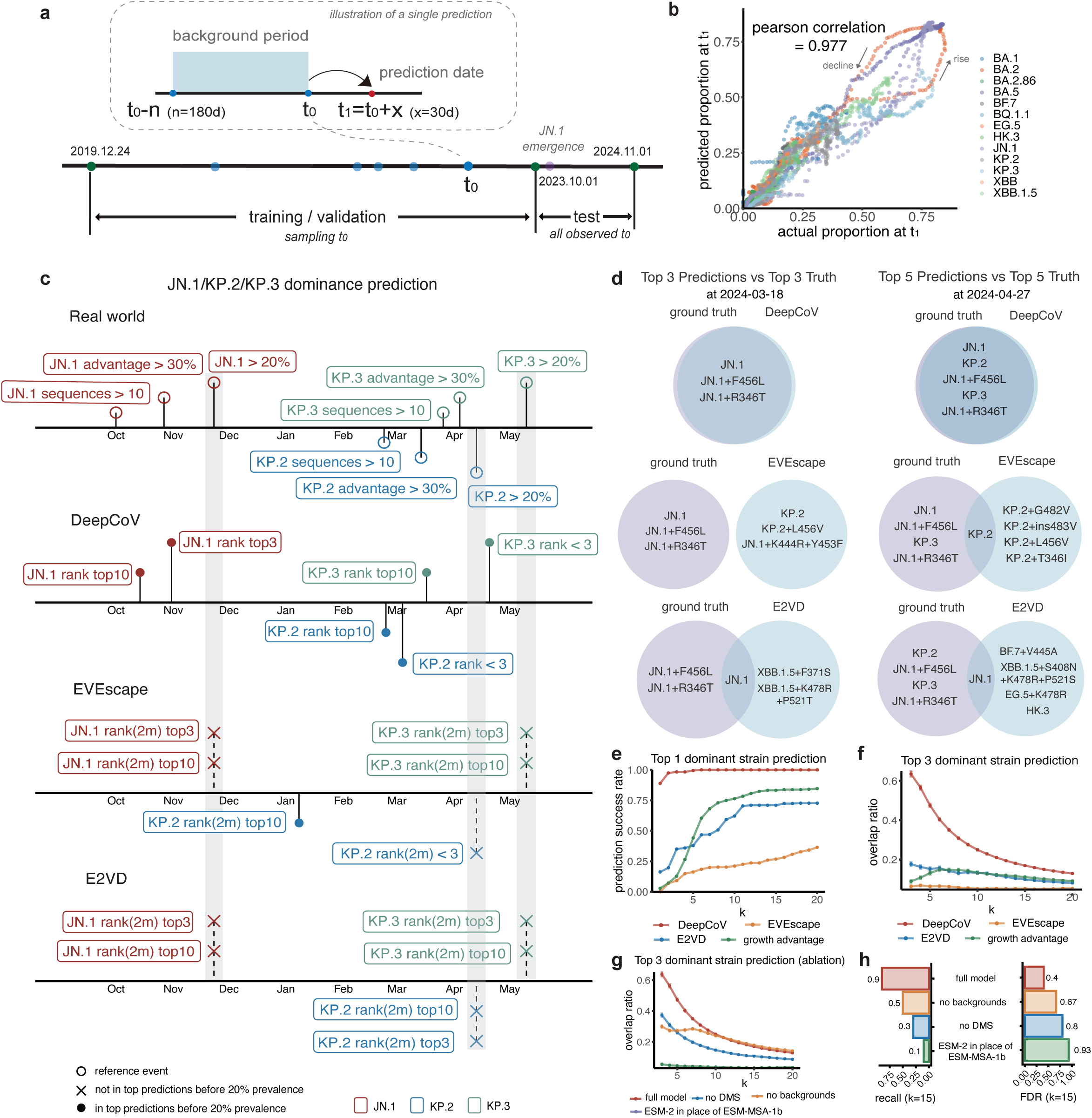
Accurate early detection of predominant strains by DeepCoV. **a**, Dataset construction. Training sequences were collected prior to October 1, 2023, including variants that exceeded 0.5% prevalence for at least one day during the follow-up period. Validation sets were generated using cluster-based sampling at a 1:10 ratio to minimize temporal data leakage. **b**, Scatter plot comparing predicted versus observed variant frequencies at the evaluation time point (t₁). Colors representing different RBD lineages. **c**, Timeline of prediction milestones for DeepCoV and baseline methods (E2VD and EVEscape), showing the performance (ranking reaching top 3 or top 10) for known dominant variants JN.1, KP.2, and KP.3. Actual emergence events, like ≥10 sequences, growth advantage > 30% and prevalence >20%, are labeled as reference points. For E2VD and EVEscape, rankings include variants with sequences appearing within a 2-month window. Solid circles connected by solid lines indicate successful predictions; dashed lines ending with a cross (×) denote prediction failures. The light grey shaded region highlights the period during which the true prevalence exceeded 20%. The inability to rank a dominant variant in the top predictions within this window is considered a failed prediction. **d**, Representative comparisons of top-k predicted and observed dominant variants at selected time points. Venn diagrams show the overlap between the predicted and observed top-k RBD variants for *k* = 3 (right; 2024-03-18, predicted 30 days in advance) and *k* = 5 (left; 2024-04-27). Predictions from DeepCoV (top row), EVEscape (middle row), and E2VD (bottom row) are compared against the corresponding observed sets. Variant labels indicate RBD lineage with key escape mutations. **e**, Top-k prediction performance over time for three methods: DeepCoV (red), EVEscape (orange), E2VD (blue) and growth advantage (green). The prediction success rate is shown for the top predicted dominant variant across varying k values (i.e., the number of predicted variants considered). **f**, Top-k prediction performance over time for three methods: DeepCoV (red), EVEscape (orange), E2VD (blue) and growth advantage (green). The jaccard index is shown for the top 3 predicted dominant variant across varying k values (i.e., the number of predicted variants considered). Error bars represent ±1 s.e.m. (standard error of the mean) across all evaluation time points. **g**, Top-k prediction performance over time for the ablated models. The jaccard index is shown for the top 3 predicted dominant variants across varying k values (i.e., the number of predicted variants considered). Error bars represent ±1 s.e.m. across all evaluation time points. **h**, FDR and recall metrics of the ablated models when k=15 for the top 10 circulating variants prediction during the JN.1 period. The top 10 true circulating variants includes JN.1, KP.2, KP.3, HK.3, JN.1+R346T, JN.1+F456L, HK.3+A475V, KP.2+G482V+K484E, KP.2+L456V+K478T, and KP.2+ins483V+K484E.

DeepCoV demonstrated high predictive capacity, evidenced by a strong correlation (Pearson’s r = 0.969) for historically dominant lineages (Fig. 2b). We systematically compared DeepCoV with conventional growth advantage fitting method and state-of-the-art deep learning models, including E2VD and EVEscape, to evaluate their performance in meeting real-world pandemic surveillance requirement^58,66^. We first assessed how early these methods could correctly prioritize the known dominant variants (JN.1, KP.2, and KP.3) among the top predicted lineages. DeepCoV uniquely identified JN.1, KP.2, and KP.3 as top dominance candidates among all the RBD sequences appeared since October 2023, well ahead of their observed dominance. In contrast, EVEscape successfully predicted only KP.2, and E2VD failed to detect any of the dominant variants (Fig. 2c).

Detailed benchmarking across different numbers of top-ranked predicted variants confirmed DeepCoV’s superior ability to identify emerging dominant lineages. To comprehensively assess predictive performance, we conducted evaluations using both time-resolved dominant variants and a fixed set of globally prevalent strains under varying *k*-candidate thresholds. Ranking-based evaluation was adopted to minimize biases introduced by raw score distributions. For the time-resolved ground truth, we evaluated two metrics across time points: the success rate of top-1 versus top-*k* prediction, and the Jaccard overlap ratio between the top-*k* predicted and top3 or top5 observed variants (Fig. 2d,e and Extended Data Fig. 2). For the static task involving a fixed set of globally prevalent strains, the objective was to rank variants with the highest overall prevalence appeared at the top. Across both evaluation schemes, our model consistently outperformed baseline methods, especially at lower *k* thresholds (Fig. 2e,f and Extended Data Fig. 2). For both top-3 versus top-3 and top-5 versus top-5 comparison, DeepCoV successfully identified all major variants; in contrast, other methods achieved at most one overlapping variant (Fig. 2d).

As for the fixed-ground-truth evaluation using the top globally prevalent strains across dates in the test set, DeepCoV also achieved notably higher recall and substantially lower false discovery rates (FDR) at top-*k* candidate thresholds below 40 (Extended Data Fig. 3a,b,d,e). For top 3 dominants prediction, DeepCoV correctly identified JN.1, KP.2, and KP.3 as the subsequently prevailing lineages at *k* = 5, with closely related subdominant variants JN.1+F456L ranked immediately below (Extended Data Fig. 3c). To further assess robustness, we relaxed the dominance criterion to include the top 10 most prevalent variants and evaluated performance at *k* = 20. Even under this broader definition, DeepCoV continued to outperform baseline methods (Extended Data Fig. 3f). Overall, its predictive accuracy remained consistent across a range of *k* values, and was particularly strong at lower *k* thresholds. This ability to ensure that true dominant variants are captured within a small set of candidate sequences has significant implications for timely vaccine design and targeted public health interventions.

The fitness scores predicted by DeepCoV have an intrinsic quantitative interpretation, allowing them to be directly mapped to real-world variant prevalence. To account for varying definitions of dominance, we assessed model sensitivity using a dynamic three-stage benchmarking framework based on surveillance data: T1 (≥10 sequences, initial emergence), T2 (>30% growth advantage over a 7-day window with ≥100 sequences), and T3 (prevalence exceeding 5–50% while maintaining >15% growth advantage). For each RBD sequences appeared since October 2023 and reach T1, we evaluated prediction success relative to T2 and T3 (Extended Data Fig. 4). Across all prevalence threshold beyond 5%, DeepCoV achieved high recall, and accuracy with low FDR. At 35% prevalence threshold, all three predictions were correct. In contrast, the commonly used growth advantage–based method yielded 63 candidate variants. These results highlight DeepCoV’s ability to accurately identify dominant variants without relying on growth advantage, while also enhancing the efficiency of growth-based methods by substantially narrowing the candidate space.

To assess the broader applicability of DeepCoV, we retrospectively evaluated its performance on earlier SARS-CoV-2 XBB lineage data, using a dataset partitioned at September 2022 and restricted to available DMS profiles from BA.1, BA.2, and BA.5 (Extended Data Fig. 5). Despite the reduced size and scope of the training set, DeepCoV achieved a strong correlation between predicted and observed prevalence (Pearson’s *r* = 0.957) and correctly identified the subsequent emergence of dominant variants XBB.1.5 and BQ.1.1, while slightly overestimating the prevalence of HV.1. These results underscore its robustness and predictive potential across distinct phases of SARS-CoV-2 evolution.

To investigate the contributions of individual model components and biological data modalities, we performed systematic ablation studies. Four model variants were constructed by selectively excluding key information: (1) immune background profiles, retaining only sequence and prevalence data; (2) DMS phenotypes, preserving sequence and immune trend inputs and replace DMS module with linear layers; and (3) evolutionary sequence context, replaced by ESM-2 embeddings to isolate the effect of evolutionary modeling. Removal of any individual module resulted in a marked decrease in predictive performance (Fig. 2g,h and Extended Data Fig. 2b,c).

We conducted the ablation experiments under both the dynamic dominant strain selection and global dominant strain selection settings, and compared the RMSE of models trained under these different conditions. Both the DMS and immune-background modules were critical for accurately detecting dominant strains, substantially reducing FDR while maintaining recall. Moreover, eliminating the proportion or ESM-MSA-1b modules rendered the model incapable of training. Together, these results highlight the essential and collaborative contributions of immune landscape dynamics, evolutionary sequence context, historical prevalence trends, and functional mutational phenotypes to the overall predictive capacity of DeepCoV.

### DeepCoV captures variant spatiotemporal dynamics

Beyond accurate early variant identification, DeepCoV effectively captures the spatiotemporal dynamics of SARS-CoV-2 variant spread, demonstrating substantial improvements over existing methodologies. We further reconstructed the evolutionary trajectories of dominant SARS-CoV-2 variants with high temporal resolution using DeepCoV. The model effectively captured the full expansion and decline cycles of major JN.1 clades, maintaining stable predictive accuracy throughout the JN.1-dominant period (Fig. 3a). DeepCoV maintained consistently high precision over time, with major strain predictions showing slightly larger but acceptable fluctuations (mean absolute error (MAE) <0.1; root mean square error (RMSE) <0.15) (Extended Data Fig. 8). Importantly, DeepCoV demonstrated sensitivity to subtle early growth signals, successfully forecasting rising of dominant variants even from low initial prevalence (<5%). Notably, despite differing from its ancestral BA.2.86 lineage by only a single RBD substitution (L455S), the rapid rise of JN.1 was correctly captured by DeepCoV as the dominant variant. This highlights DeepCoV’s capacity to distinguish variants with minimal genetic differences but markedly divergent epidemiological trajectories, underscoring its sensitivity to functionally meaningful mutations. In addition to major lineages, the model faithfully reconstructed the growth trajectories of subdominant variants such as JN.1+F456L and JN.1+R346T, as well as high-growth advantage but ultimately low-prevalence lineages including JN.1+K403R and JN.1+N417K (Extended Data Fig. 8). DeepCoV also showed high specificity in handling non-dominant variants, with predicted peak prevalence consistently remaining below 3%. Moreover, even when the training set is restricted to data prior to the emergence of XBB, the model also successfully reconstructed the growth trajectories of all major variants in the test set. Notably, even when trained exclusively on pre-XBB data, DeepCoV successfully forecasted the emergence of JN.1 sublineages (Extended Data Fig. 5b). These findings underscore the model’s utility for monitoring fine-scale viral evolution and guiding timely public health responses, including vaccine strain selection.

**Figure 3.**
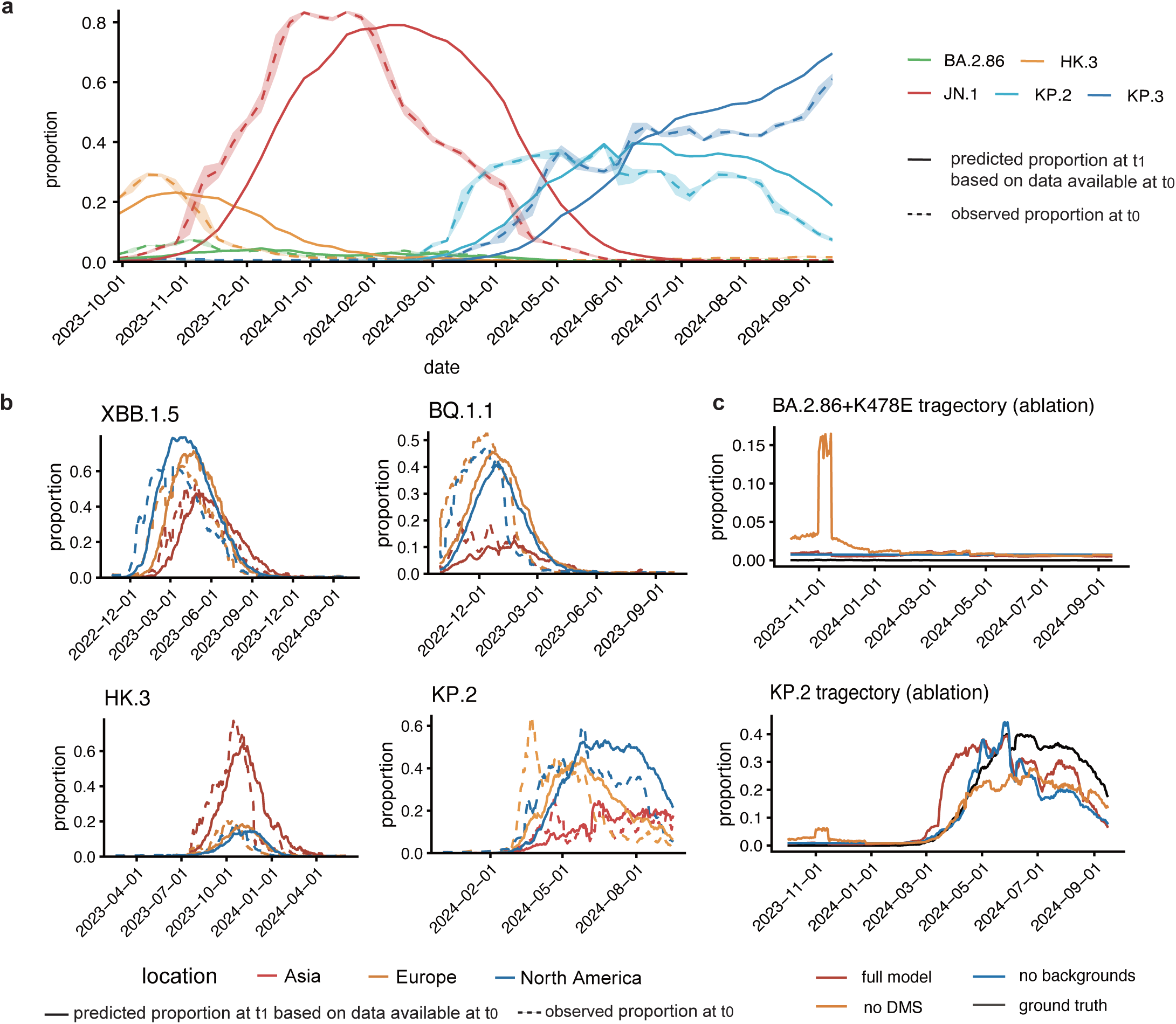
DeepCoV captures temporal dynamics and geographic variation in SARS-CoV-2 spread. **a**, Growth trajectory reconstruction. Weekly aggregated temporal dynamics comparing predicted (dashed lines, proportion at s predicted from t_0_) versus actual prevalence (solid lines, observed at t_0_). Shaded regions represent mean ± standard deviation. **b**, Regional prevalence patterns across Asia, Europe and North America. Predicted (dashed lines) and observed (solid lines) variant proportions were aggregated weekly. **c**. Ablation study on growth trajectory reconstruction for BA.2.86 + K478E and KP.2. The black solid line indicates the full model and the dashed lines represents the ablated variants.

By intrinsically incorporating regional immune landscape variations, DeepCoV successfully captures geographically distinct transmission patterns. The model accurately reconstructed intercontinental divergence patterns, clearly capturing the sequential emergence of KP.2 from Europe to North America and subsequently Asia (Fig. 3d). It also correctly identified elevated prevalence of variants such as BQ.1.1 and XBB in Western populations relative to Asia, while highlighting the regional dominance of HK.3 in Asia, reflecting regional differences in pandemic responses and immune imprinting. Although other models may account for temporal factors, DeepCoV uniquely combines spatiotemporal resolution with proactive forecasting and quantitatively validated accuracy, offering improved interpretability in complex epidemiological settings.

Ablated models were further assessed on growth trajectory reconstruction (Fig. 3e). The *no DMS* variant erroneously overestimated the growth advantage of BA.2.86+K478E, underscoring the essential role of the DMS module in mitigating false positives. Moreover, removing any single module eliminated the early prediction of KP.2, indicating that complementary signals from multiple modules are required to support the model’s overall performance.

To achieve finer period forecasting, we developed a continuous prediction model employing a LSTM network capable of forecasting variant trajectories over subsequent 1 to 60-day windows (Extended Data Fig. 8). This approach maintained robust predictive performance with low RMSE, comparable to the baseline model, although a slight decline in accuracy was observed over extended forecasting intervals (Extended Data Fig. 8b). Importantly, it was also capable of anticipating the future trajectory of dominant variants over a sustained period, even from early time points (Extended Data Fig. 8c). Whereas the 30-day model focuses on short-term qualitative dominance prediction, the continuous model characterizes long-term prevalence dynamics, pinpointing when variant expansion accelerates or decelerates. This framework enhanced DeepCoV’s suitability for real-time surveillance and proactive public health planning.

### In silico mutational hotspot scanning

Leveraging DeepCoV’s ability to capture evolutionary dynamics, we conducted analysis to in silico identify mutational hotspots within the SARS-CoV-2 RBD, aiming to understand the driving forces behind immune escape mutations during convergent evolution. We computationally generated all possible single-site RBD mutants for representative convergent lineages, including JN.1 and XBB variants, and applied models trained on temporally matched datasets from the respective JN.1 or XBB eras. By predicting time-resolved evolutionary scores for each mutation, we dynamically mapped site-specific evolutionary pressure and identified candidate hotspots likely to contribute to future adaptation (Fig. 4a).

**Figure 4.**
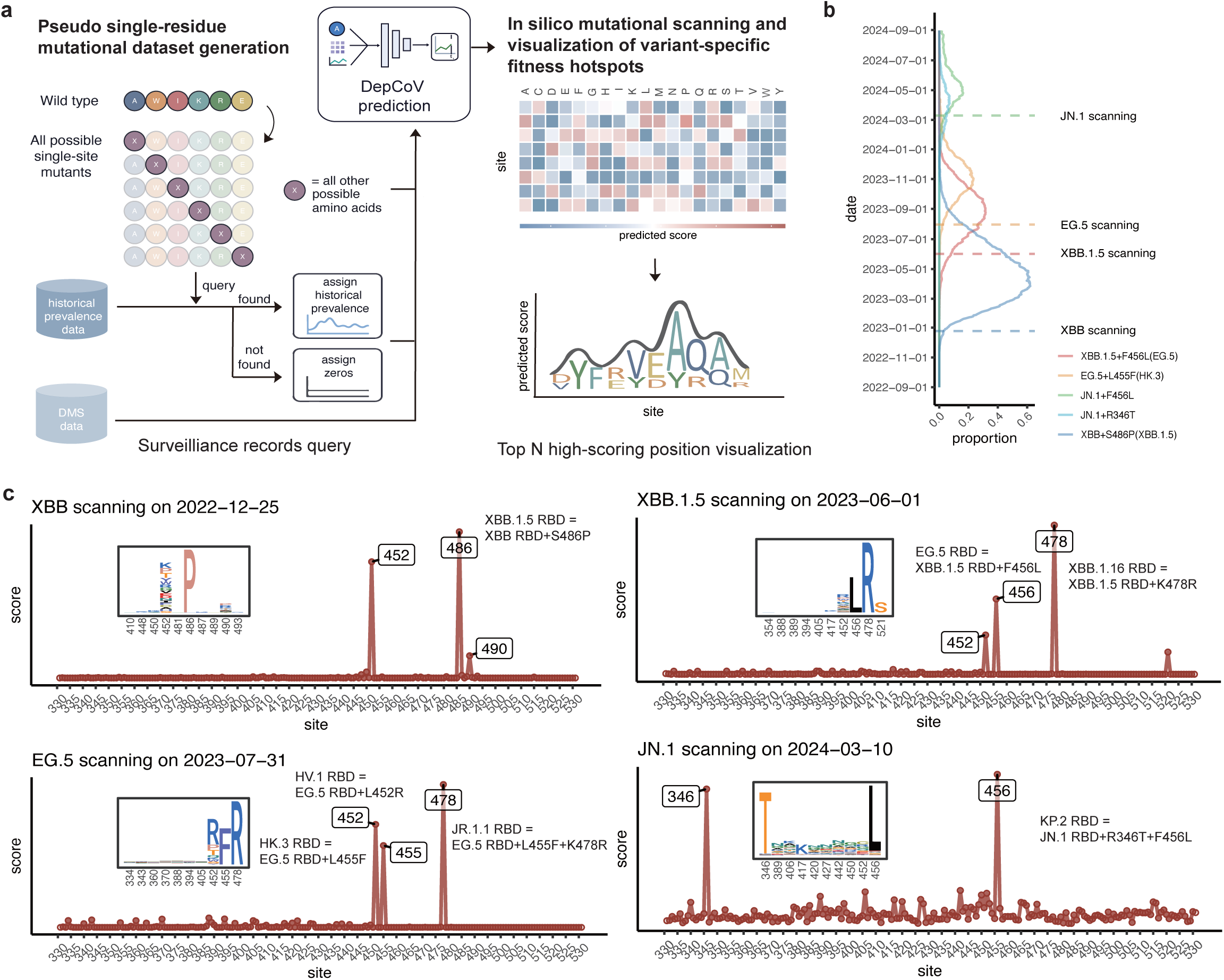
Dynamic mutational hotspot scanning in silico. **a,** Schematic of the mutational scanning workflow. Pseudo single-residue mutational datasets were generated and evaluated using the predictive model to estimate evolutionary potential for each mutation. **b,** Temporal prevalence dynamics of variants containing convergent mutational hotspots. Vertical dashed lines indicate selected timepoints for in silico scanning. **c,** Predicted mutation preference landscapes for JN.1, XBB, XBB.1.5 and EG.5 prior to the onset of convergent evolution.

Our analysis successfully identified mutational hotspots at residues R346T and F456L in JN.1, which later became defining mutations in emerging strains such as KP.2 before their prevalence achieving 5% (Fig. 4c)^41^. As expected, site like 403, previously overestimated by DMS-based assays, did not exhibit notable predicted evolutionary potential. Similarly, DeepCoV accurately identified key mutational hotspots associated with subsequent variant dominance, including the early prediction of the S486P substitution, followed by F456L and L455F mutations within the XBB lineage^25,84,85^. These predictions accurately forecasted the sequential “FLip” mutation wave, reflecting real-world evolutionary trajectories. Importantly, DeepCoV identified these mutation patterns significantly ahead of widespread detection; for instance, the S486P hotspot could be predicted before XBB.1.5 (XBB with the S486P mutation) became detectable in global sequencing data (Fig. 4b). Subsequently, the emergence of residues 455 and the combination 455+456 mutations was identified in early evolutionary stages of variants like EG.5 and HK.3, aligning with structural insights suggesting compensatory functional interplay between these residues. Our findings demonstrate that DeepCoV effectively captures intrinsic residue-level drivers of evolutionary convergence. By combining temporal modeling of mutation phenotypes with sequence-based predictions, our approach mirrors the functional resolution provided by DMS experiments but with added temporal insights into mutation dynamics.

We further assessed the contribution of individual modules to JN.1 mutational hotspot detection and forecasting of future evolutionary trajectories (Extended Data Fig. 9). All ablated model variants exhibited a pronounced loss of hotspot discrimination, with only the F456L mutation detected in the *no DMS* model—likely reflecting prior selection signals from lineages such as XBB, where F456L conferred marked escape potential. Collectively, these complementary modules act synergistically to support reliable prediction of future mutational trends.

### DeepCoV generalizes to future SARS-CoV-2 evolution

Recently, increased immune pressure on the spike protein’s N-terminal domain (NTD), which facilitates viral entry, has led to elevated mutational activity, establishing it as a secondary hotspot of adaptive evolution^86–89^. Variants such as XEC (T22N/F59S) and KP.3.1.1 (S31del) exemplify this trend, reflecting the evolving selective pressures shaping SARS-CoV-2’s immune escape mechanisms^73,81,90–93^. To capture the evolutionary shift, we extended DeepCoV to be trained on the full spike protein sequences, recalculating prevalence metrics based on unique spike clusters while preserving the original model architecture (Extended Data Fig. 10). Considering that most DMS measurements outside the RBD region are unavailable, which could interfere with model training, the updated model excluded the DMS module. The effect of removing this component could be partially compensated by the expanded number of Spike sequences incorporated into the training data.

The refined approach maintained excellent prediction accuracy and consistently low FDR for dominant variants, while recall briefly decreased at intermediate then subsequently recovering at higher thresholds for widely circulating variants. Moreover, the updated model robustly reconstructed the evolutionary trajectories of complex variants such as KP.2.3 (KP.2+S31del+H146Q) and KP.3.1.1 (KP.3+S31del), demonstrating strong generalizability. In addition, in silico mutational scanning of the NTD successfully identified S31 deletions as potential immune escape mutations, which had been suggested to enhance immune escape through allosteric modulation of RBD–antibody interactions mediated by additional NTD glycosylation^91^. These results underscore DeepCoV’s robust capability to generalize to previously underrepresented structural domains and highlight its utility in modeling future evolutionary adaptations.

We updated the test dataset till May 2025 to evaluate DeepCoV’s performance against recently emergent strains, including LP.8 and KP.3+A435S whose peak prevalence exceeded 10% globally. The model maintained strong predictive accuracy (Pearson’s *r* = 0.968; Fig. 5b), accurately forecasting the emergence of LP.8 and continuing to perform well on expanded lineages such as JN.1, KP.2, and KP.3. Retrospective in silico mutational scanning of KP.3 and LF.7 revealed early detection of future-dominant mutations (Fig. 5a,d). Notably, A435S was identified as a prominent hotspot approximately one month before the widespread emergence of KP.3+A435S, while residue 475, harbored by later emerging LF.7.2.1 strain, was similarly highlighted during pre-emergence scans. We further tested the model’s ability to resolve geographic prevalence differences by evaluating its prediction of NB.1.8.1 spatial dynamics^5^. As shown in Fig. 5c, DeepCoV accurately identified the disproportionately high prevalence of NB.1.8.1 in Asia relative to other regions. Together, these findings demonstrate DeepCoV’s continued ability to anticipate the emergence and geographic distribution of newly arising variants and mutational hotspots, even beyond its original training horizon.

**Figure 5.**
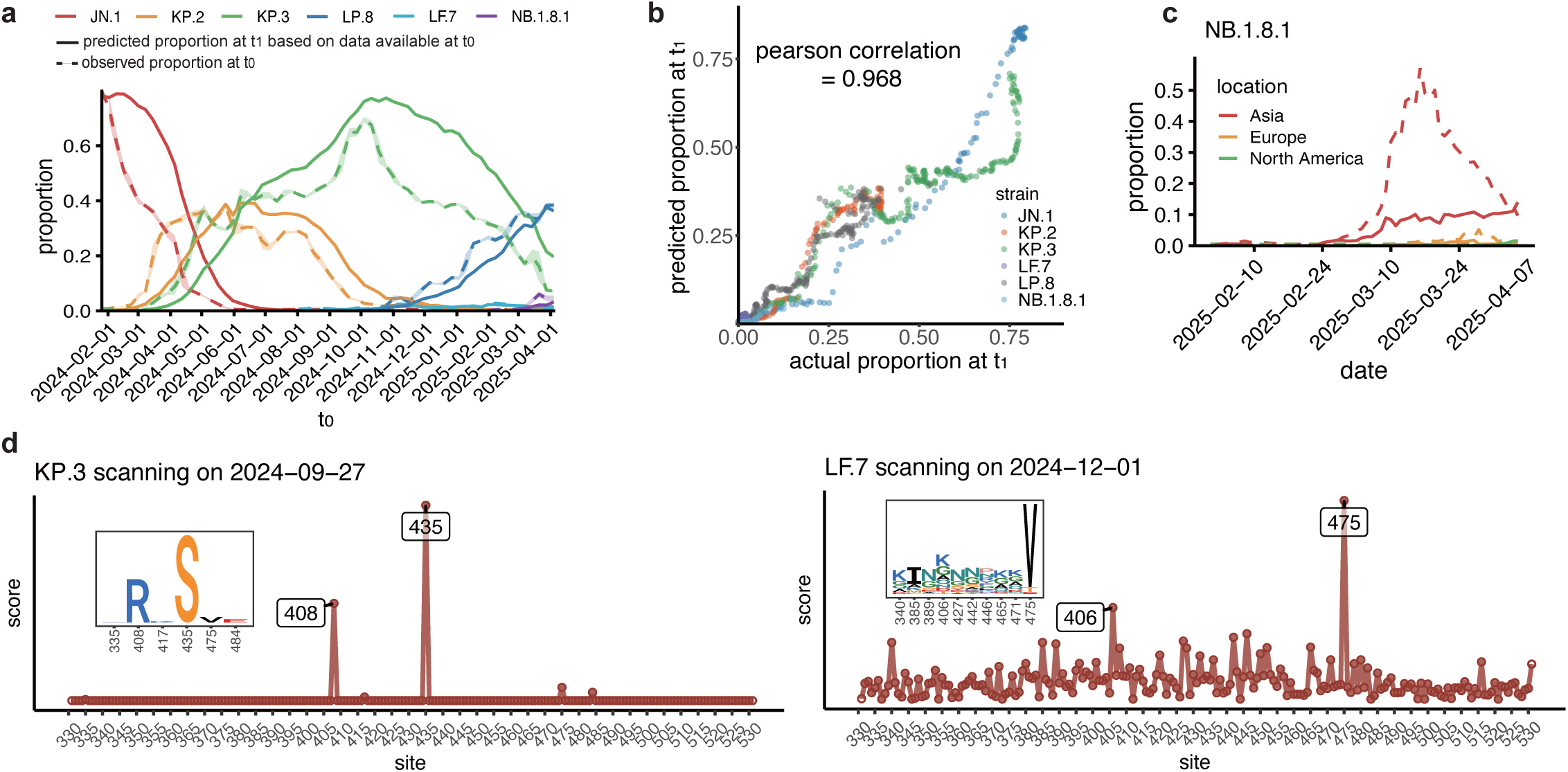
Updating SARS-CoV-2 variant data for future predictions. **a**, Growth trajectory reconstruction using the renewed test set. Weekly aggregated predictions (dashed lines, t_1_ inferred from t_0_) are compared with observed prevalence (solid lines, measured at t_0_). Shaded regions represent mean ± s.d for days in a week. **b**, Pearson’s correlation coefficient of predicted versus observed variant frequencies at time, for each strain prior to reaching its peak prevalence. Colors representing different RBD lineages. Each point represents a variant, colored by lineage. **c**, Predicted regional differences in NB.1.8.1 prevalence across Asia, Europe and North America. Predicted (dashed) and observed (solid) proportions are aggregated weekly. **d**, Prevalence of site-specific mutations in KP.3 and LF.7 prior to the emergence of convergent evolution, identifying early mutational hotspots.

## Discussion

One of the major challenges in SARS-CoV-2 surveillance and vaccine design lies in both the timely identification of emerging dominant variants after their emergence and the anticipation of high-risk lineages before they arise. To address this, we developed DeepCoV, a computational framework that integrates key evolutionary drivers—viral MSA patterns, mutational phenotypes from DMS, and epidemiological data reflecting historical immune pressures—to effectively predict SARS-CoV-2 variant prevalence. DeepCoV employs the ESM-MSA-1b to capture evolutionary constraints from sequence alignments, integrates DMS-derived phenotypic data through Transformer-based modules, and models temporal epidemiological trends using LSTM networks. By integrating sequence constraints, DMS-derived mutational effects, and temporal prevalence trends, and by formulating the training objective to emphasize early prediction of dominant strains, DeepCoV achieves markedly lower FDR and reliably identifies dominant variants ahead of conventional surveillance methods. Moreover, it captures global and regional variant prevalence trajectories and successfully predicts evolutionary mutational hotspots in different dominant strain eras, serving as a robust and scalable tool for real-time surveillance of emerging viral populations. Ultimately, DeepCoV offers an effective early warning system that enhances public health preparedness by supporting timely policy decisions, optimizing vaccine strategies, and guiding surveillance efforts.

Despite its strengths, DeepCoV has several limitations. The scarcity of comprehensive DMS data for certain protein domains—particularly regions outside the RBD—together with inherent sequencing errors in epidemiological datasets, may affect predictive performance. Moreover, epistatic effects are not explicitly modeled, and extending computational predictions to incorporate combinatorial mutations from DMS data could improve the representation of these interactions^47,49,53,70,94^. Future refinements might also integrate additional data modalities, such as structural protein information, to further enhance predictive accuracy and generalizability^76,89^. Meanwhile, although DeepCoV indirectly reflects population-level immunity through prevalence dynamics, it does not yet capture the full complexity of the evolving immune landscape. Future *in silico* virus–immunity co-evolution models that jointly learn viral antigenicity and host immune adaptation may offer a more mechanistic understanding of immune-driven viral evolution. Finally, the reliability of DeepCoV’s predictions is influenced by the breadth and representativeness of available sequence prevalence data, highlighting the value of continued global genomic surveillance.

In conclusion, by integrating the comprehensive epidemiological and DMS datasets with frontier deep-learning-based protein language models, DeepCoV unifies evolutionary, functional, and epidemiological insights to build a reliable platform for the identification and prediction of prevalent SARS-CoV-2 strains. The model could be retrained and utilized in other fast-evolving epidemic viruses with enough datasets, such as influenza and RSV, once corresponding DMS datasets become available. Collectively, these innovations establish DeepCoV as a powerful tool for global health preparedness, enabling proactive responses to emerging infectious threats and informing timely vaccine and surveillance strategies.

## Acknowledgments

We are grateful to Jesse D. Bloom and Tyler N. Starr for making their DMS libraries and data publicly available. We extend our thanks to Jing Wang, Zheng Bian, and all supporting technicians for their contributions in dataset preparation and preprocessing. We also appreciate the scientific community for their ongoing surveillance of SARS-CoV-2 variants and valuable discussions. F.J. is supported by the Boya Postdoctoral Fellowship Program of Peking University and the Postdoctoral Fellowship Program of China Postdoctoral Science Foundation under Grant Number GZC20250980. Y.C. is financially supported by National Natural Science Foundation of China (32222030), the Ministry of Science and Technology of China (2021D0102) and Changping Laboratory (2025D0401).

## Author Contributions

Y.C. designed and supervised the study. S.Y., F.J., and Y.C. wrote the manuscript. F.J., X.L., S.Y., and J.L. conceived the computational model architecture. S.Y. and X.L. performed model training and optimization. S.Y. and J.L. conducted data preprocessing and curated the experimental datasets. S.Y. executed comprehensive benchmark evaluations and performed temporal-geospatial analyses.

## Declaration of Interests

Y.C. is the inventor of the provisional patent applications for BD series antibodies, which includes BD55-5514 (SA55). Y.C. is the founder of Singlomics Biopharmaceuticals. Other authors declare no competing interests.

## Methods

### Data and preprocessing

We obtained SARS-CoV-2 sequences with collection/submit dates from GISAID and retained human spike entries after de-duplication and MAFFT alignment^95,96^. Quality filters required spike length >1,230 aa, ≤10 non-standard residues, and ambiguity-free RBD; lineage-specific insertions (e.g., BA.1 ins214:EPE; BA.2.86 ins16:MPLF) were preserved. Unique RBDs were clustered and renamed relative to parental lineages for downstream modeling.

### Spatiotemporal dataset construction

For each representative region (global, North America, Europe, Asia, USA, UK, Japan), daily counts of unique RBDs were computed and 7-day smoothed. Training/validation data used t₀ snapshots with 16 top circulating background clusters in the preceding 180 days. At each reference date t_0_, the prediction target was the variant’s relative prevalence 30 days later (t_1_ = t_0_+30d). Ground-truth labels were retained only when the cumulative number of sequences in the t_1_ evaluation window was ≥100. We stratified t₀ across five pandemic phases, limited post-2023 analyses to global counts due to coverage decline, and split train/validation (90:10) by sequence to avoid leakage.

### Deep mutational scanning features

We standardized DMS datasets covering entry efficiency, ACE2 binding, expression, and serum/monoclonal-antibody escape from public and internal sources, aligning them to spike and indexing by antigen, feature, site, mutant, and (where applicable) antibody. Temporal masking ensured only features available prior to t₀ were used; antibody escape was re-clustered into 56 epitope groups; per-sequence vectors were produced by scanning one-hot sequences against aligned DMS tensors.

### Model overview

DeepCoV integrates (i) frozen ESM-MSA-1b embeddings of the target and contemporaneous background sequences; (ii) a background-ratio encoder that summarizes 180-day variant frequency histories; and (iii) a DMS encoder gathering sequence phenotypes. Sequence and background signals are fused via an axial attention module (row/column transformers) and a transformer encoder that sequentially incorporates DMS features. Outputs are future variant proportions at a single horizon (t₁ = t₀+30 d). Additional details are provided in the Supplementary Methods.

### Training and objective

Models were trained with AdamW (learning rate 10^−4^, weight decay 10^−2^) with 300-step warm-up and mixed precision under the PyTorch framework on NVIDIA A100 GPUs. The loss is a log-transformed, sample-weighted MSE with (i) a validity mask based on t₁ coverage (≥100 isolates) and (ii) labelled proportion-dependent weights to mitigate class imbalance; for the continuous model, an additional per-day Gaussian weight (μ=30, σ=10) emphasizes informative t_1_ horizons. Early stopping was applied as specified.

### Baselines and benchmarking

**Growth advantage.** For each RBD cluster, daily frequency f(t) is fit by a logistic curve to estimate growth rate *a*. Growth advantage is defined as GA=e^a × g^ − 1, with generation time g=7 days; 95% CIs are reported.

**EVEscape.** For each candidate RBD, mutations are computed relative to a reference and permutation EVEscape scores are aggregated to yield a composite sequence-level score, which is used to rank variants in the evaluation window.

**E2VD.** We re-trained E2VD on an ESM-2 backbone and combined three sub-modules—ACE2 binding, expression, and antibody escape—using an asymmetric scheme that penalizes decreases in expression/ACE2 (below functional thresholds) and rewards increases in escape (above a permissive cutoff). The final score is the sum of exponentiated deviations from empirically defined thresholds.

**Temporal/Ranking evaluations.** We (i) track monthly dominant variants by taking the mode of day-wise winners per month, harmonizing method outputs (prevalence for DeepCoV/GA; score ranks for other models) and simulating surveillance lag for DeepCoV; (ii) quantify timeliness as the earliest date a well-known dominants enters top-N (e.g., 3 or 10); and (iii) run a dynamic top-k comparison over time. For multiple-truth settings, we report the Jaccard index

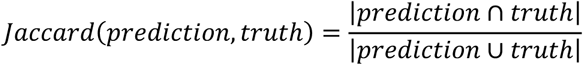

and for single-truth settings we report prediction success rate (exact cover). (iv) run a static top-k comparison for well-known dominants.

### Generalization

**Updated JN.1-era setting.** We retain the main model trained on data before 1 Oct 2023 and **extend** testing to 16 May 2025, with a lineage map that renames unique RBDs relative to parental references. A major-strain panel (JN.1, KP.2, KP.3, LF.7, LP.8, NB.1.8.1) is used for targeted trajectory evaluation.

**Spike variant.** To satisfy ESM-MSA-1b input limits, spike inputs are truncated to the first 1,023 amino acids (a minor biological compromise for C-terminal tail) while keeping all other processing consistent.

**XBB-era variant.** To accommodate reduced training data, we decreased the depth of the MSA-proportion fusion transformer from three to two layers; other components were unchanged.

**Continuous model.** The output head is replaced with an LSTM to emit daily proportions; the loss includes a per-day Gaussian weight to emphasize mid-range horizons applied uniformly across samples together with a validity mask for insufficient coverage. (Full loss and masking details in the Supplementary Information.)

### In silico mutational scanning

We generated pseudo single-amino-acid libraries on reference backbones (e.g., XBB.1.5, JN.1, KP.2/3, LF.7) for RBD (331–531) and NTD (14–305; including deletions), scored mutants with era-matched models, and summarized site-level fitness by averaging positive, residue-normalized contributions; top positions were visualized via smoothed profiles and sequence logos.

### Ablation studies

We quantify contributions of major modules via: (i) **Sequence encoder swap** (ESM-MSA-1b → ESM-2-150M) with other components fixed; (ii) **No-DMS** (replace DMS encoder with two feed-forward layers to match dimensionality); and (iii) **No background strains** (encode only the target sequence and its 180-day prevalence; aggregate histories via LSTM and a transformer feature aggregator).

## Extended Data Figures

**Extended Data Fig.1.**
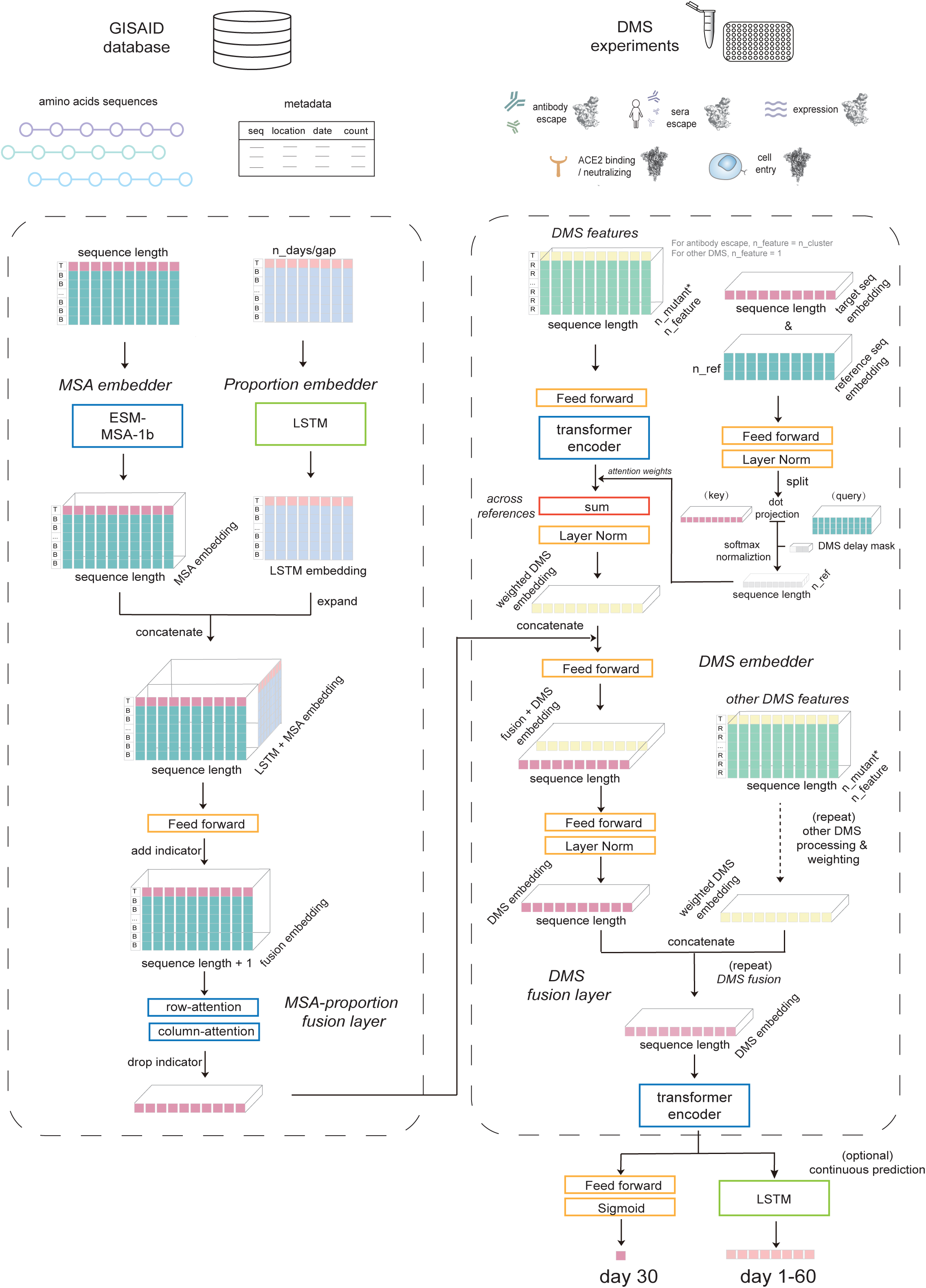
Detailed schematic of model architecture.

**Extended Data Fig.2.**
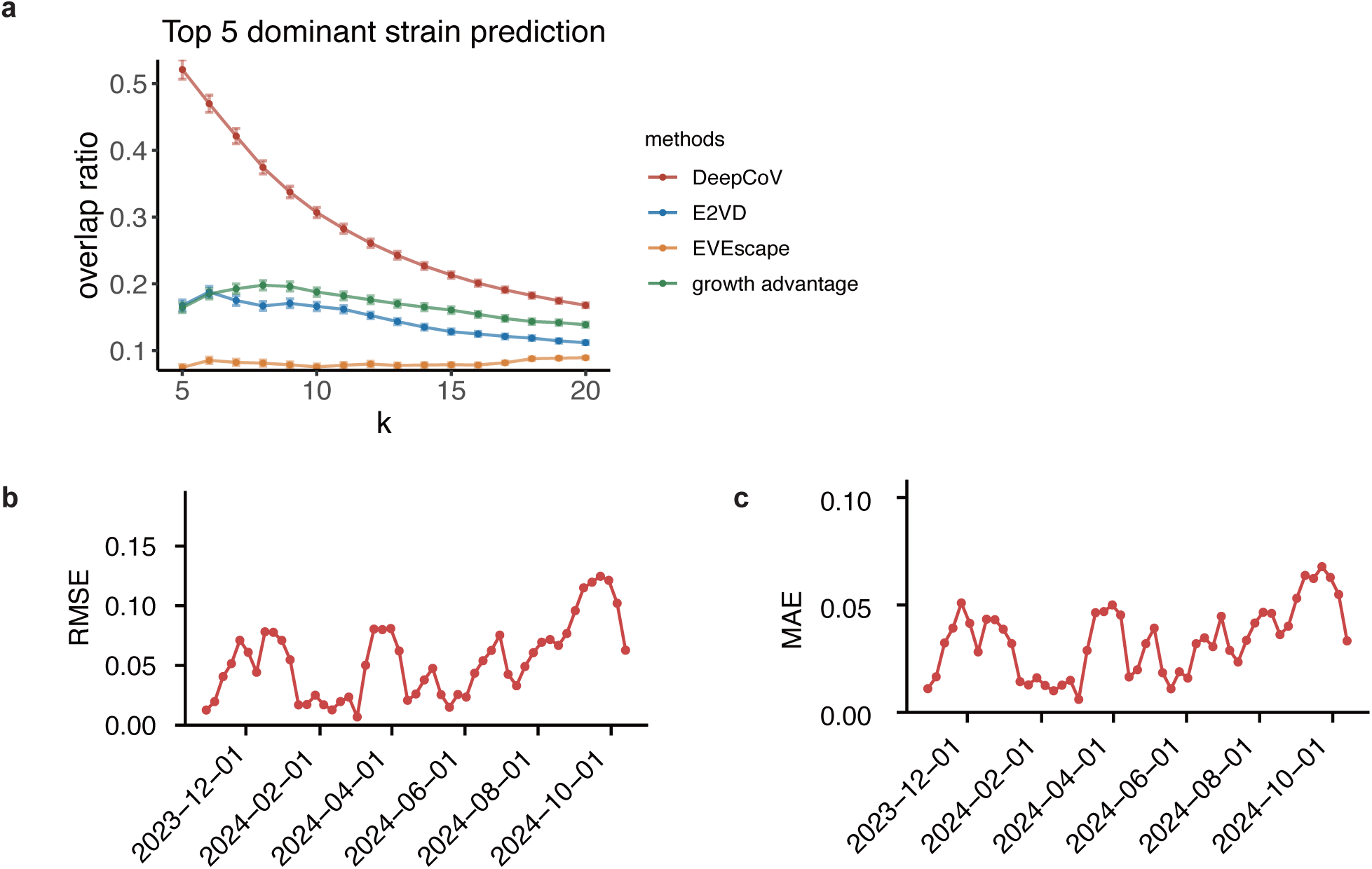
Comprehensive evaluation of dominant variant prediction accuracy and temporal dynamics. **a,** Mean Jaccard overlap ratios between predicted and observed sets of dominant SARS-CoV-2 RBD variants across varying values of *k* (number of top-ranked predictions considered) for top 5 ground-truth variants; **b**, Root mean squared error (RMSE) and **c**, mean absolute error (MAE) evaluated monthly for dominant variants after 1 October 2023 (HK.3, BA.2.86, JN.1, KP.2, KP.3).

**Extended Data Fig.3.**
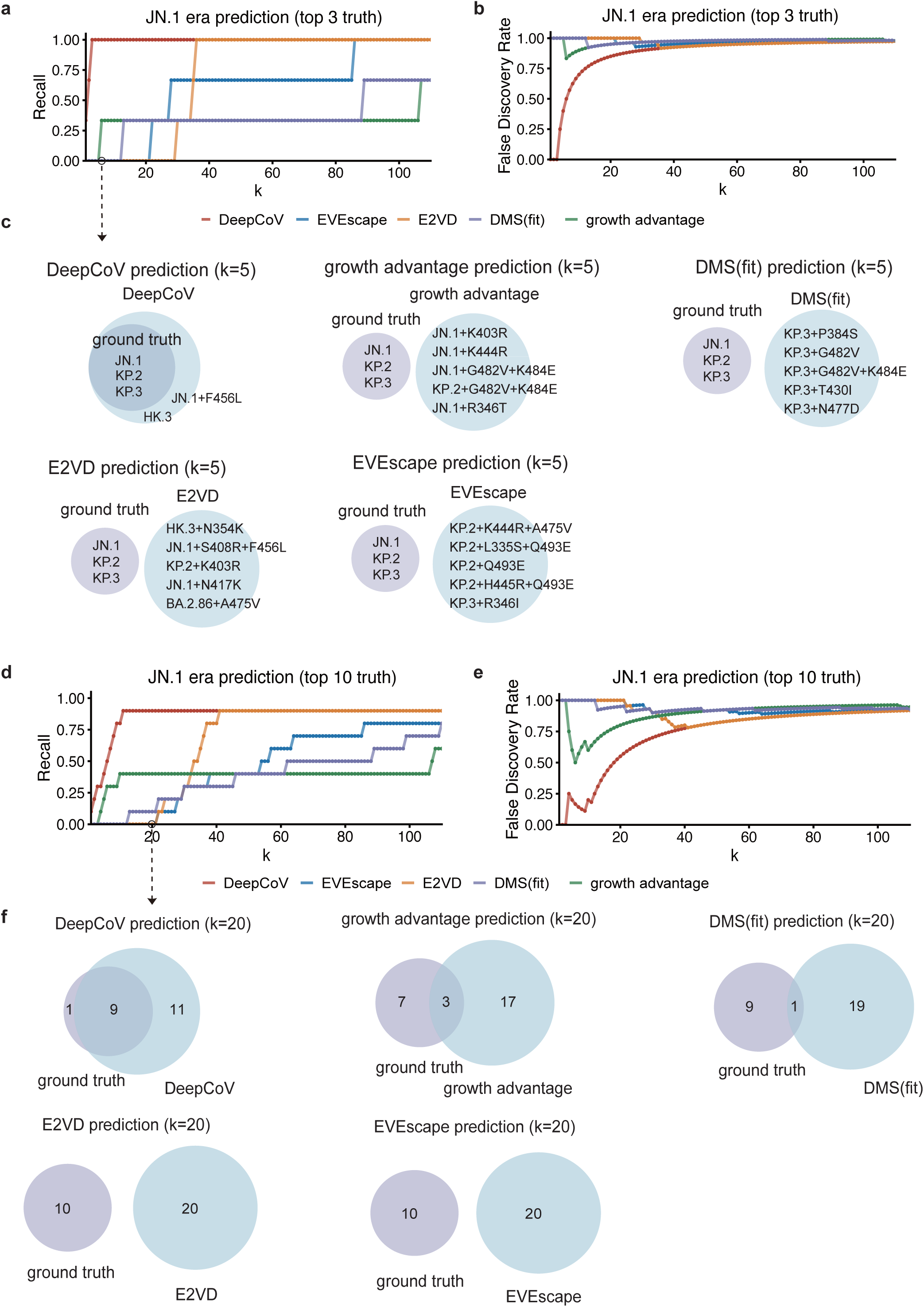
Evaluation of model performance in predicting the top 10 dominant variants during the JN.1 era. **a–b**, Performance of five methods—DeepCoV, EVEscape, E2VD, DMS(fit), and growth advantage—in recovering true dominant variants across increasing top-*k* prediction thresholds. **a**, Recall computed against three major circulating variants (JN.1, KP.2, and KP.3). **b**, Corresponding false discovery rate (FDR). **c**, Venn diagrams showing predicted (purple) and ground truth (blue) dominant variants at *k = 5* (top) for each method. **d–e**, Extended recall (**d**) and FDR (**e**) performance across *k* for the top 10 circulating variants during the JN.1 period, defined as: JN.1, KP.2, KP.3, HK.3, JN.1+R346T, JN.1+F456L, HK.3+A475V, KP.2+G482V+K484E, KP.2+L456V+K478T, and KP.2+ins483V+K484E. **f**, Venn diagrams comparing predicted versus observed dominant variants at *k = 20*.

**Extended Data Fig.4.**
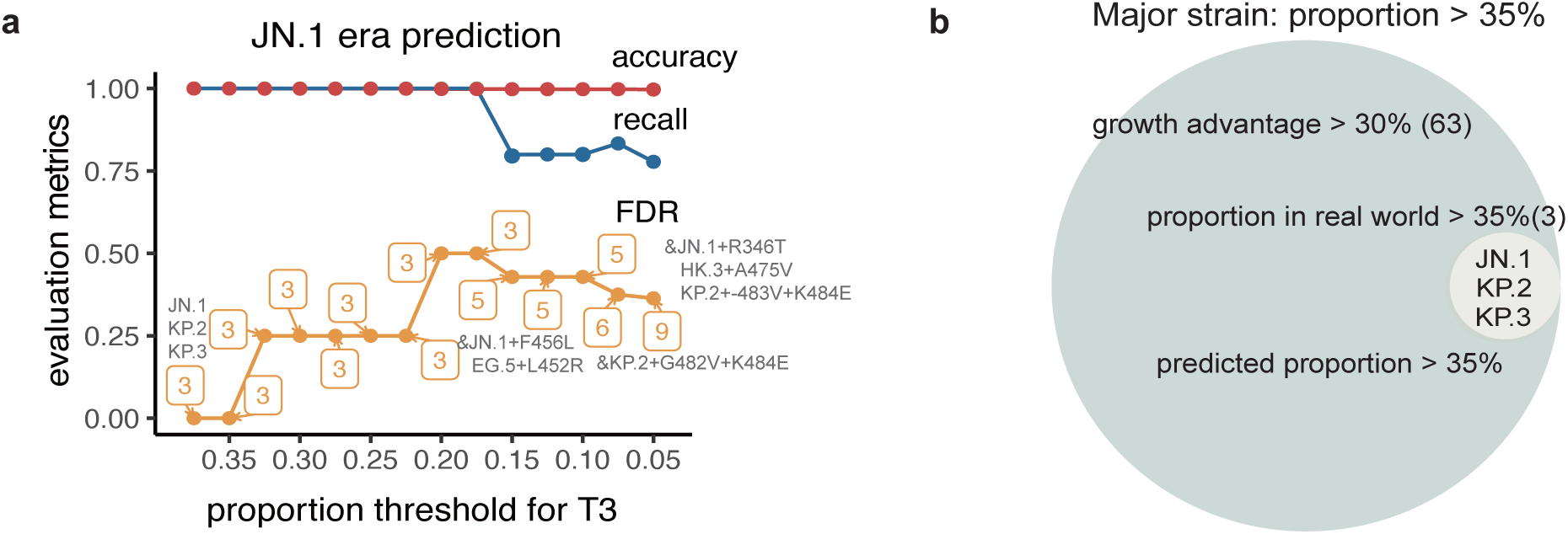
Dynamic evaluation of model performance across variant prevalence thresholds during the JN.1 era. **a**, Model performance metrics—accuracy, recall, and FDR—evaluated under varying prevalence thresholds used to define dominant variants. The FDR curve is annotated with the number of actual dominant variants for each prevalence threshold, along with the corresponding true variants added for each proportion threshold. **b,** Comparison of predicted versus actual dominant variants under threshold definitions of >35% variant prevalence. The number of strains identified by each methods are labeled in parentheses.

**Extended Data Fig.5.**
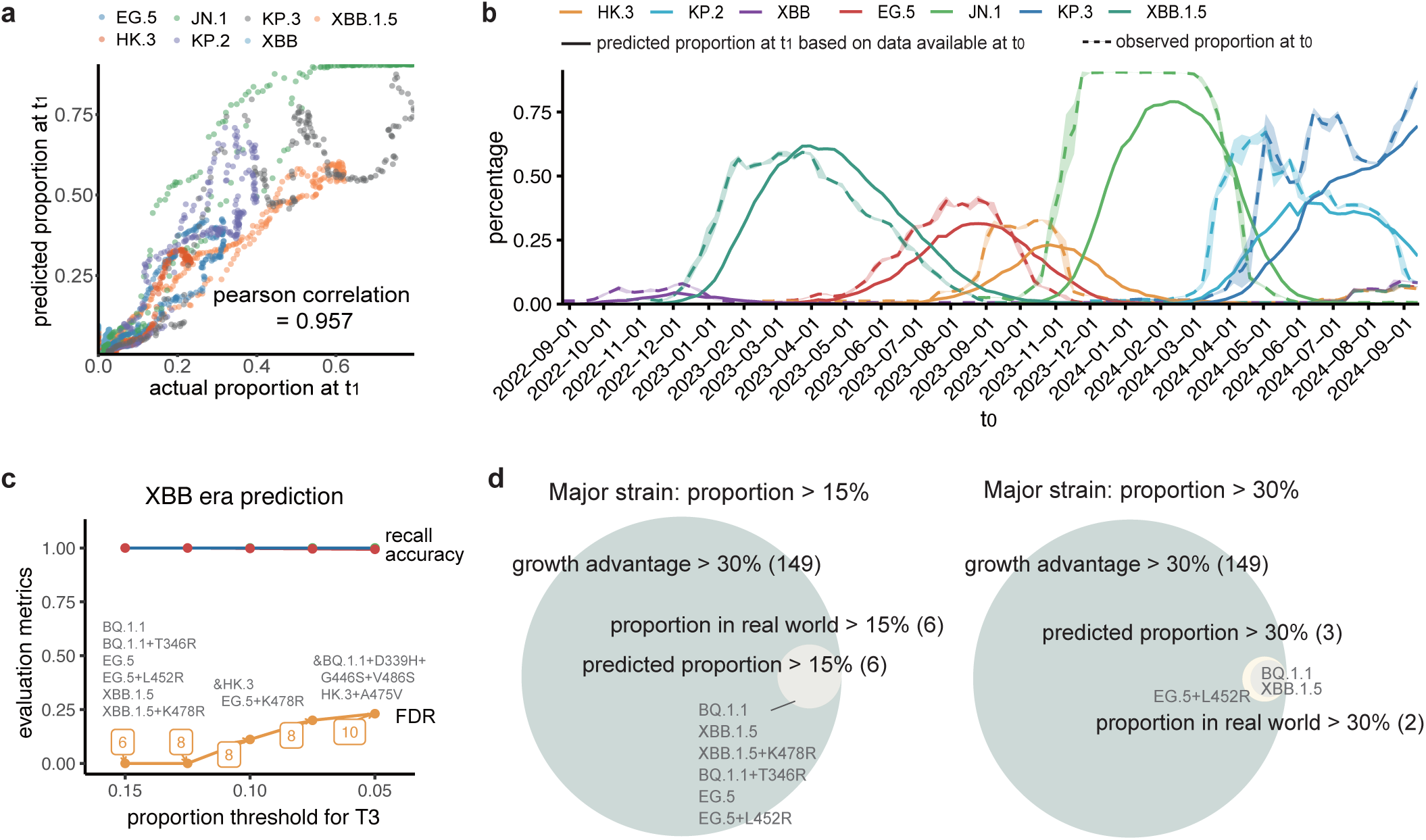
DeepCoV performance on XBB-era variants. **a**, Correlation between predicted and observed variant frequencies at evaluation time (t₁) using a model trained on spike sequences post-XBB emergence, for each strain prior to reaching its peak prevalence. Colors representing different RBD lineages. Each point denotes a variant, colored by lineage; Pearson’s correlation coefficient (r) is indicated. **b**, Growth trajectory reconstruction of dominant variants after 1 September 2022. Weekly aggregated predictions (dashed lines, t₁ inferred from t₀) are compared with observed prevalence (solid lines, measured at t₀). Shaded areas represent mean ± s.d. across each week. **c**, Dynamic assessment of model performance across varying definitions of dominant variants based on prevalence thresholds. Accuracy, FDR, and recall are reported under each threshold setting. The FDR curve is annotated with the number of actual dominant variants for each prevalence threshold, along with the corresponding true variants added for each proportion threshold. **d,** Comparison of predicted versus actual dominant variants under threshold definitions of >15% and >30% variant prevalence. The number of strains identified by each methods are labeled in parentheses.

**Extended Data Fig.6.**
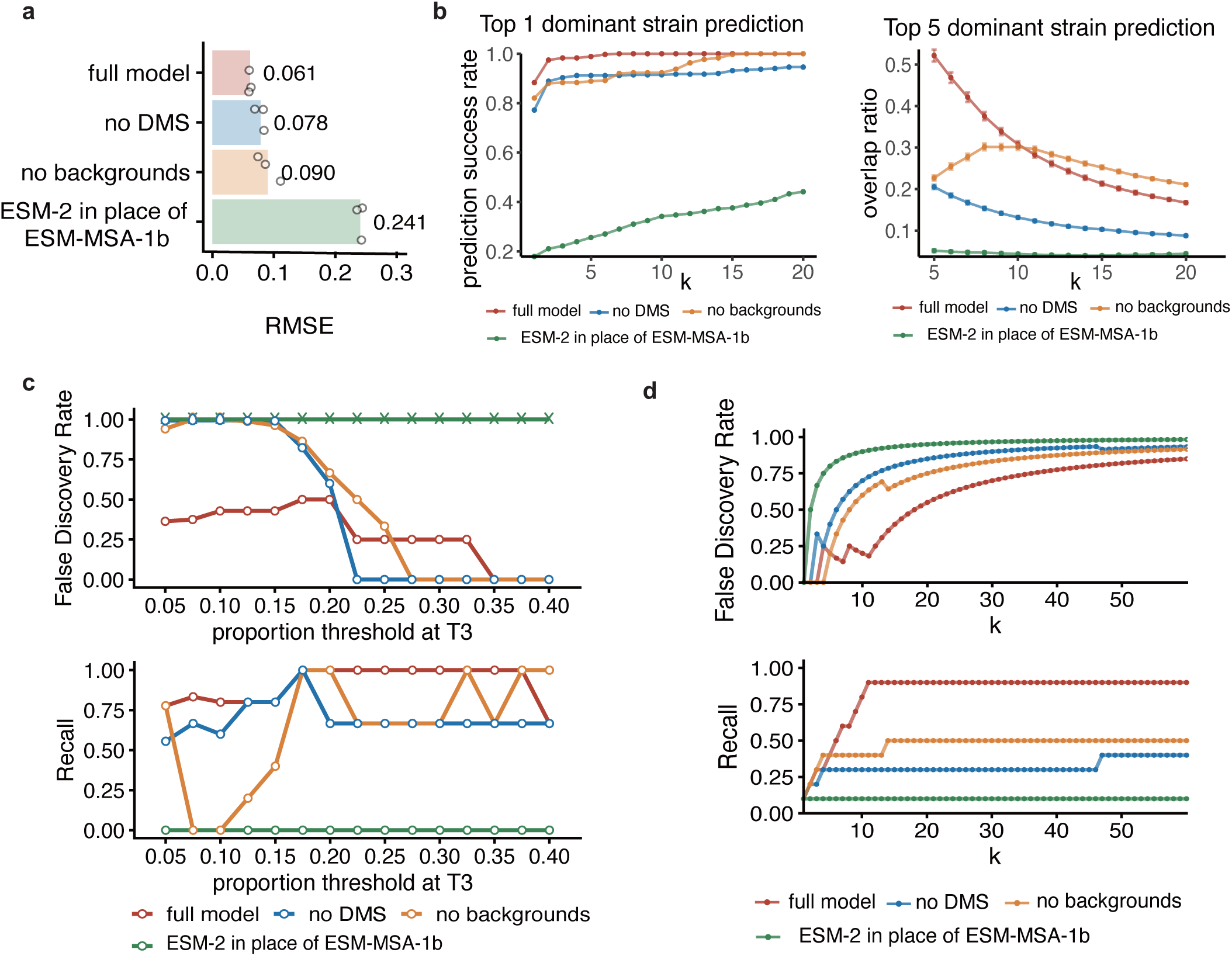
Ablation study evaluating the contribution of individual components to DeepCoV performance. **a,** Comparative RMSE across the full model and three ablated models: (i) removal of immune background features, (ii) exclusion of DMS data, and (iii) replacement of evolutionary features with ESM-2 embeddings. **b**, Top-k prediction performance over time for the ablated models. Success rate is reported for the top 1, and jaccard index is reported for the top 5 predicted dominant variants across varying *k* values (i.e., the number of predicted variants considered). Error bars represent ±1 s.e.m. across all evaluation time points. **c,** Predictive metrics (false discovery rate and recall) stratified by variant prevalence thresholds across ablated models. **d,** Performance of ablated model in recovering true dominant variants across increasing top-*k* prediction thresholds, evaluated using recall, FDR and accuracy computed against top ten major circulating variants (JN.1, KP.2, KP.3, HK.3, JN.1+R346T, JN.1+F456L, HK.3+A475V, KP.2+G482V+K484E, KP.2+L456V+K478T, and KP.2+ins483V+K484E).

**Extended Data Fig.7.**
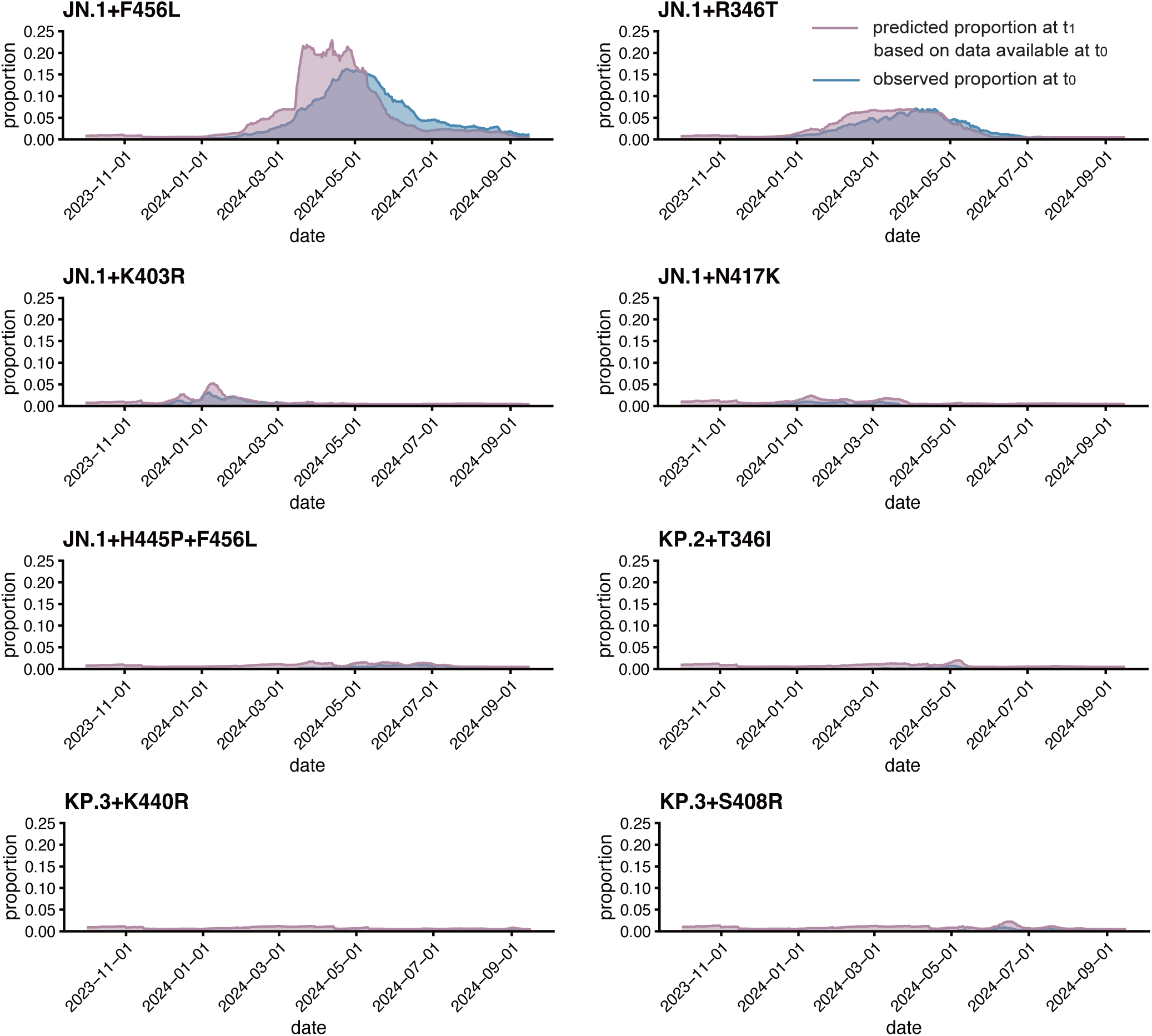
Model performance in detecting minor SARS-CoV-2 variants. Growth trajectory reconstruction for subdominant and low-prevalence variants. Weekly aggregated predictions (lines colored purple, t_1_ inferred from t_0_) are compared with observed prevalence (lines colored blue, measured at t_0_).

**Extended Data Fig.8.**
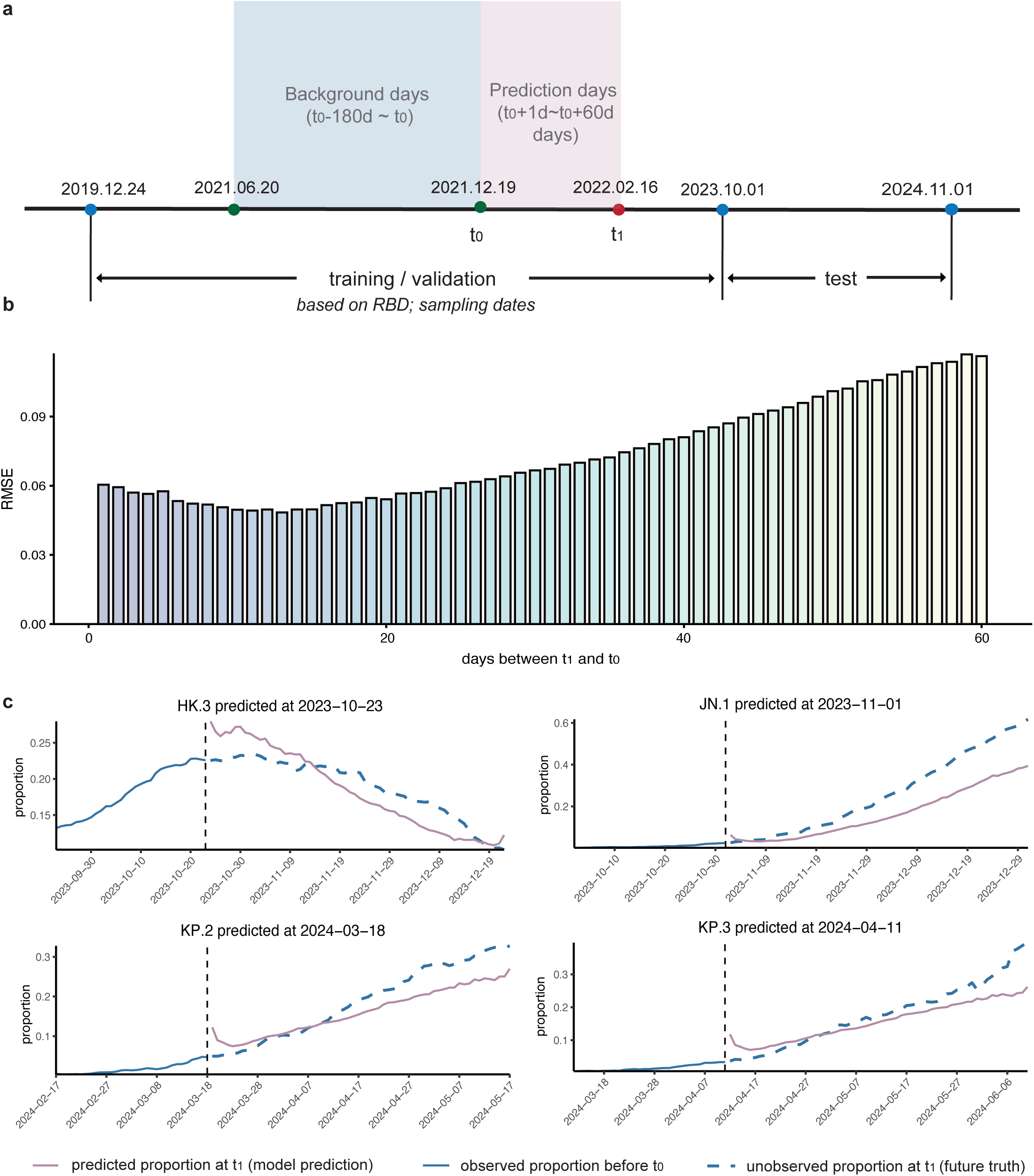
Longitudinal performance evaluation of predictive model. **a**, The dataset composition of model continuous prediction. For each t_0_, the future prevalence for 1-60 days (t_1_) later are predicted. **b**, Evaluation of model performance on the major lineage test set across varying forecasting horizons, with a fixed t_0_ and incrementally increasing t_1_. **c**, Early forecasting of future prevalence trajectories (1–60 days ahead) for dominant variants. Purple lines show model-predicted prevalence at future time points (t_1_, t_0_+1 to t_0_+60) using data available at t_0_; blue solid lines show observed prevalence before t_0_; blue dashed lines show observed prevalence at the corresponding future time points (t_1_).

**Extended Data Fig.9.**
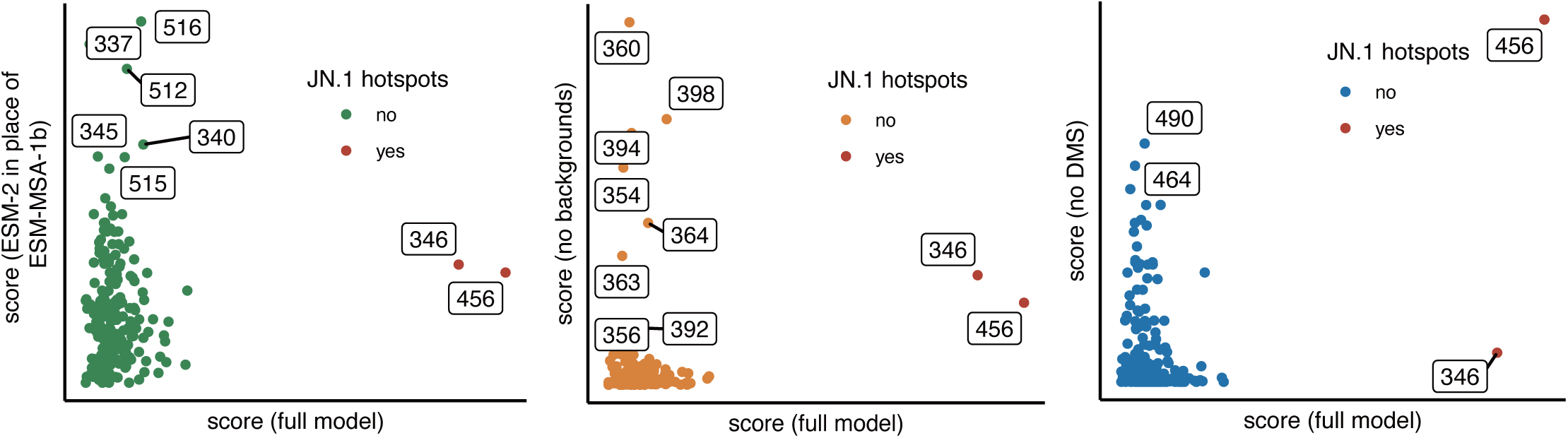
Ablation on in silico mutational hotspots scanning. Comparison of the full and ablated models in detecting JN.1 mutational hotspots. True hotspots (positions 346 and 456) are highlighted in red. In silico deep mutational scanning of JN.1 single-point mutants at 20 March 2024 illustrates the predicted fitness landscapes under each ablation setting.

**Extended Data Fig.10.**
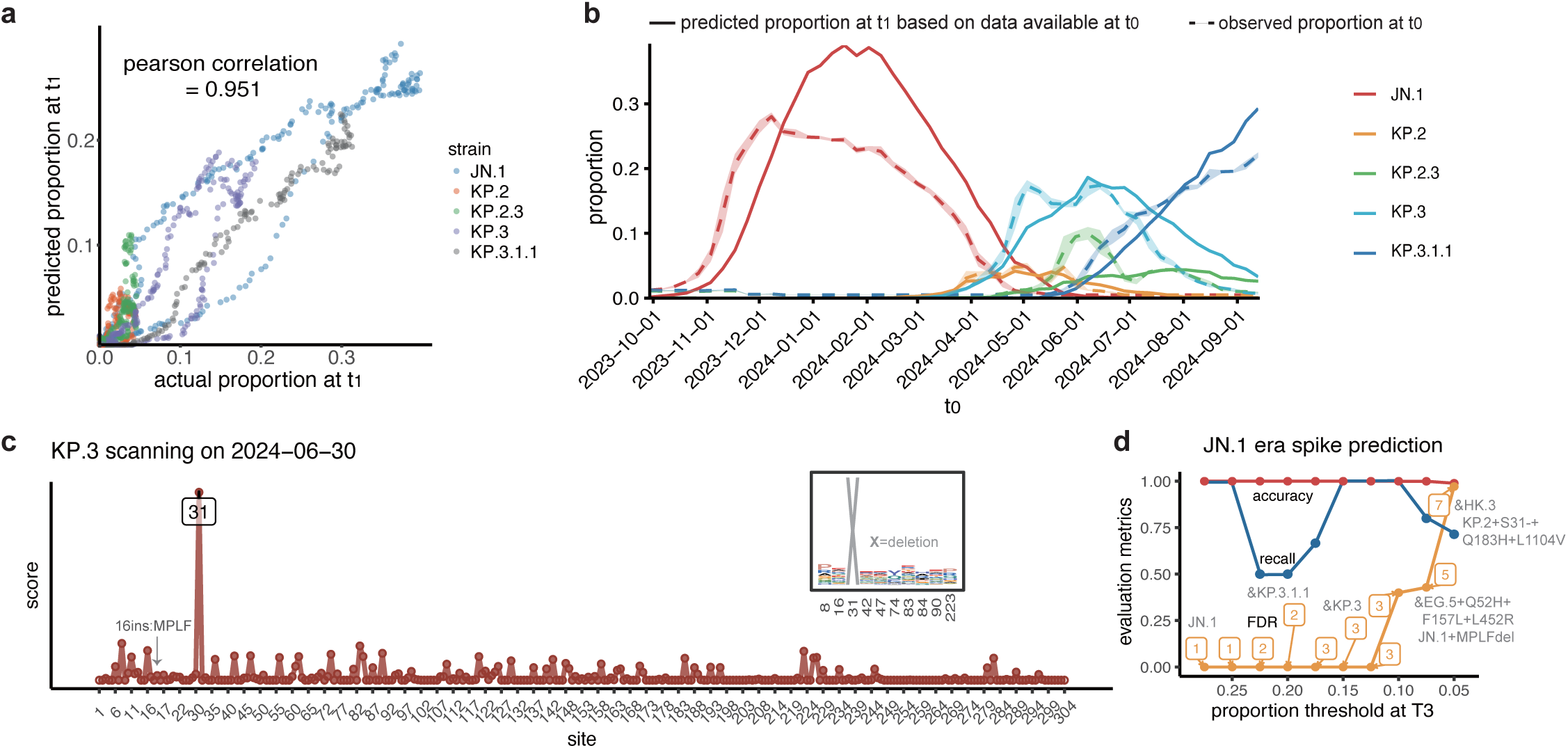
DeepCoV performance of model trained on spike protein. **a**, Pearson’s correlation coefficient (r) of predicted versus observed variant frequencies at time t_1_ using model trained on SARS-CoV-2 spike. Each point represents a variant, colored by lineage. **b**, Growth trajectory reconstruction using the renewed test set. Weekly aggregated predictions (dashed lines, t_1_ inferred from t_0_) are compared with observed prevalence (solid lines, measured at t_0_). Shaded regions represent mean ± s.d for days in a week. **c**, Prevalence of site-specific mutations and deletions in KP.3 prior to the emergence of convergent evolution, identifying early mutational hotspots. “X” denotes deletion mutations. **d**, Dynamic assessment of spike-based model performance across varying definitions of dominant variants based on prevalence thresholds. Accuracy, recall, and FDR are reported under each threshold setting. The number of actual dominant variants for each prevalence thresholds are labeled.

