## Supplementary Methods for "Predicting SARS-CoV-2 evolution dynamics with spatiotemporal resolution by DMS-empowered protein language model"

Data preprocessing

SARS-CoV-2 sequence and epidemiological data preprocessing

SARS-CoV-2 viral sequences and related location and date metadata were downloaded from GISAID^1^. For preprocessing, sequences were first filtered based on the metadata to include only original human spike proteins with complete collection dates and submitted dates (YYYY-MM-DD format). After deduplication, sequences were aligned to the reference spike proteins using MAFFT^2^, and further filtered to >1230 residues, ≤10 non-standard residues, and quality-controlled RBD regions are extracted without ambiguous residues. The common insertions ins214:EPE harbored by BA.1 and ins16:MPLF harbored by BA.2.86 sub-strains are retained in the multiple sequence alignments. Following quality control, 28,837 unique high-quality RBD sequences and 743,737 unique high-quality spike sequences were used for subsequent procedure. For each unique RBD sequences, cluster index and cluster name are assigned for further usage. Unique RBD are renamed according to their relative mutations relative to their parental lineage references: WT, Alpha, Beta, Delta, Gamma, Eta, BA.1, BA.2, BA.5, BF.7, BQ.1.1, XBB, XBB.1.5, EG.5, HK.3, BA.2.86, JN.1, KP.2 and KP.3.

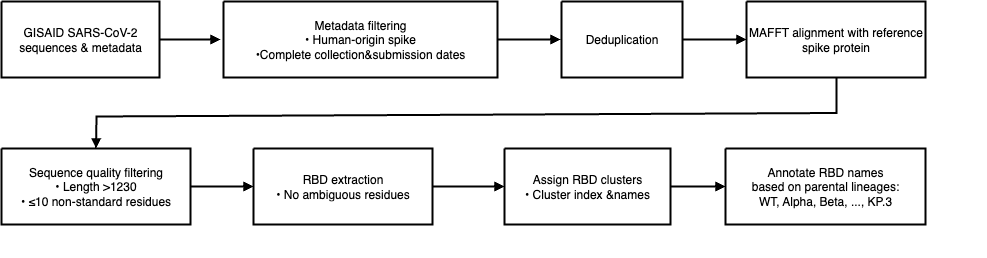

Figure S1 ｜Workflow of SARS-CoV-2 sequence data preprocessing

Sequence datasets construction and spatiotemporal stratification

We constructed sequence dataset annotated with spatiotemporal metadata as follows. We compute the count of each unique RBD and the total counts in each region per days. To ensure data stability and reliability, we restricted the analysis to the global total and six representative regions: North America, Europe, Asia, USA, United Kingdom, and Japan, while excluding unique RBD with fewer than 30 total sequences locally. To mitigate high-frequency noise on sequencing day-level variability, we performed 7-day window smoothing on the sequence count. For efficient indexing and aggregation, we constructed enumerators to map each collection date, country, and continent into index spaces, allowing matrix-based count operations.

We construct spatiotemporal stratified training and validation datasets for variant prevalence modeling as follows. For each spatiotemporal (location, sequence, t₀) triplets, background clusters in the past 180 days (t₀ -180 ~ t₀) are used to indicate the immunological pressure. Each candidate (location, sequence) pair was evaluated across all possible t₀ dates using the following inclusion criteria. At time t₀, the number of distinct background clusters circulating within the preceding 180-day window must be ≥16, ensuring sufficient immunological pressure complexity observation. For a given t₀, the model predicts variant growth on 30 days later. To ensure the reliability of prediction targets, the cumulative isolate count during the t₁ interval was required to exceed 100. Data prior to October 1, 2023, were used for the training and validation sets, while data collected afterward were reserved for the test set. To balance training data quality with comprehensive test set coverage, we applied stricter filtering and sampling criteria to the training data, while retaining the test set data as completely as possible. To balance the number of positive and negative samples in the training set and enable the model to better learn the characteristics of major strains, we required that at least one day within the t₁ interval exhibit a target cluster ratio above 0.5%. To ensure coverage across the pandemic timeline, t_0_ samples are evenly drawn from five time bins: [2020.02.01–2021.07.01), [2021.07.01–2022.04.01), [2022.04.01–2022.12.01), [2022.12.01–2023.05.01), [2023-05.01–2023.10.01). Given the progressive decline in sequencing volumes across regions following 2023, especially after WHO declaration of the end of the pandemic, only global-level sequence counts were included after 2023.01.01 to maintain data quality standards. The data were further split into training and validation sets (90:10 ratio) by stratified random sampling on the (location, sequence) level to prevent data leakage. This yielded 73,081 items for the training and validation sets.

Sequence datasets construction and spatiotemporal stratification

To evaluate the predictive performance of our model on emerging SARS-CoV-2 variants, we construct two types of test sets: a comprehensive full test set and a curated major-strain-specific test set including HK.3, BA.2.86, JN.1, KP.2 and KP.3 RBDs. Both sets are based on observations collected after October 1, 2023. The full test set includes all candidate variants that emerged globally after the training cutoff date. There were a total of 741,463 entries in the full test set. Using a curated timeline of variant emergence, we assigned each major strain a reference start date based on its early documented isolation before burst. The full test set emphasizes comprehensive evaluation of overall model performance across a wide range of candidate variants, whereas the major test set specifically assesses the model’s ability to forecast the trajectories of high-priority, globally prevalent lineages and allows for detailed, trajectory-level validation of model predictions across variants of significant public health relevance.

Preprocessing of deep mutation scanning features

To integrate functional mutational data into the DeepCoV framework, we curated and standardized multiple DMS datasets from public repositories and internal lab sources. We first collected four major classes of DMS datasets, including: (i) Spike S protein-mediated entry efficiency and ACE2 binding affinity (BA.2 and XBB.1.5) from Bloom lab^3^; (ii) RBD expression and ACE2 binding data from Starr et al.^4^; (iii) serum and monoclonal antibody escape profiles including neutralization escape data from XBB.1.5 and Delta^3^; and (iv) large-scale monoclonal antibody escape data from Cao Lab^5-9^. These datasets were retrieved from published GitHub repositories or internal lab directories. Raw DMS data are formatted into consistent tabular structures and converted into structured arrays aligned with the reference Spike protein sequence for downstream model input. Each DMS array was indexed along four to five axes depending on the dataset: antigen (variant background), feature (phenotype), site (aligned residue index), mutant amino acid, and optionally, antibody identity for antibody escape array. Missing values were filled with NaNs.

To avoid data leakage and enable dynamic, time-aware inference, we indexed DMS features by their antigen source and recorded their earliest availability dates. During training, we applied temporal masking to ensure only features experimentally available prior to the target prediction date (t₀) were used. For antibody escape features, we aggregated epitope-level escape scores by antibody cluster and averaged within each group to mitigate sampling noise. We re-clustered the antibody escape profiles into 56 distinct groups to obtain a more fine-grained representation of the underlying epitope landscape. Only antibodies with prior immunological exposure (i.e., sampling time ≤ t₀) were retained for analysis. Each input sequence was one-hot encoded and scanned against the aligned DMS matrix to produce a position-specific, feature-aware vector representation.

Model architecture

The overall architecture of DeepCoV is shown in Extended Data Fig.1, and its aim is to predict the proportion in future of certain RBD or Spike. To represent the dynamic immune pressures, MSA of target and top-circulating strains at time and the corresponding sequences count for certain date and location, are collected from GISAID database. Mutation phenotype of evasion (antibody escape, sera escape), fitness (ACE2 binding, Spike mediate entry) as well as expression from Deep mutation scanning are generated by Cao et al and Bloom et al, that incorporates constraints from.

First，we adopt pretrained ESM-MSA-1b with frozen parameters to extract amino-acid-level embeddings of RBD or Spike of SARS-CoV-2 target and representative variants. The 3-day-gap-level proportion of variants generated from GISAID, with location information implicitly represented, are fed into LSTM. Subsequently, the MSA embedding and .and proportion embedding are concatenated together and processed by the self-trained DeepCoV axial-attention module^10^. In this module, a three-layer row-wise and column-wise transformer is adopted to build local dependencies of the residues and variants. The refined embedding of target are then split for further DMS information integration. As for DMS data, we express the sequence-level score of antibody escape for each epitope by max pooling on the site dimension and reference dimension, and mean across sites and references for other DMS features. To go a step further, the MSA-proportion embedding, concatenated with the DMS features one by one, flows into transformer encoder module for feature coupling. The purpose of using a transformer here is to emphasize the impact of protein-level embedding. At the final step, linear layers are applied to generate the proportion for certain day and LSTM for certain period considering the time dependence.

Sequence Encoding Module

To represent protein sequences effectively, we employ state-of-the-art protein language models that capture evolutionary constraints and contextual amino acid relationships. We support two pre-trained models:

$$\begin{aligned} E_{seq}=\text{SeqEncoder}\left( \boldsymbol{X} \right)\in\mathbb{R}^{B\times\left( 1+N_{bg} \right)\times L\times D}\#\left( 1 \right) \end{aligned}$$

where $B$ is batch size, $N_{bg}$ is the number of background sequences, $L$ is sequence length, and $D$ is embedding dimension. The main model employs the ESM-MSA-1b as the sequence encoder to exploit multiple sequence alignment information, while ablation variants optionally substitute ESM-2 to assess the impact of evolutionary modeling on downstream performance.

Background Ratio Encoder

The background ratio encoder processes temporal variant frequency data to capture evolutionary dynamics. This component enables the model to learn patterns in how variants rise and fall in prevalence over time. The encoder employs a hierarchical LSTM architecture that processes time series data of variant frequencies:

$$\begin{aligned} \boldsymbol{h}_{t},\boldsymbol{c}_{t}=\text{LSTM}\left( \boldsymbol{x}_{t},\boldsymbol{h}_{t-1},c_{t-1} \right)\#\left( 2 \right) \end{aligned}$$

where $\boldsymbol{x}_{\boldsymbol{t}}\in\mathbb{R}^{B\times N_{bg}\times1}$ represents background ratios at time $t$, and $\boldsymbol{h}_{t},\boldsymbol{c}_{t}$ are hidden and cell states.

To effectively integrate temporal patterns with sequence information, we implement an attention-based fusion mechanism. This approach allows the model to selectively focus on relevant aspects of both sequence and temporal data:

$$\begin{aligned} E_{{seq}_{proj}}=\text{Linear}\left( \text{Mean}\left( E_{seq},\text{dim}=2 \right) \right)\#\left( 3 \right) \end{aligned}$$

$$\begin{aligned} \boldsymbol{A}_{\boldsymbol{query}},A_{attn}=\text{MultiheadAttention}\left( h_{last},E_{{seq}_{proj}},h_{last} \right)\boldsymbol{\#}\left( 4 \right) \end{aligned}$$

where $h_{last}$ serves as the final hidden state of the LSTM and functions as the query; $E_{{seq}_{proj}}$ represents the projected sequence encoding acting as the key, while $h_{last}$ simultaneously serves as the value. The multi-head attention mechanism computes similarity scores between queries and keys, then performs a weighted summation of values using these scores to produce $\boldsymbol{A}_{\boldsymbol{query}}$, with $A_{attn}$ containing the attention weights. The attention mechanism computes similarity scores between the LSTM's final hidden state and projected sequence embeddings, allowing the model to focus on sequence features most relevant to temporal dynamics. We implement a gating mechanism to control information flow between temporal and sequence features:

$$\begin{aligned} \boldsymbol{g}=\sigma\left( \text{Linear}\left( \left[ \boldsymbol{h}_{\boldsymbol{last}};A_{query} \right] \right) \right)\boldsymbol{\#}\left( 5 \right) \end{aligned}$$

$$\begin{aligned} E_{bg}=\boldsymbol{g}\odot\boldsymbol{h}_{\boldsymbol{last}}+\left( 1-\boldsymbol{g} \right)\odot A_{query}\#\left( 6 \right) \end{aligned}$$

where $\boldsymbol{h}_{\boldsymbol{last}}$ is the final hidden state of the LSTM, $A_{query}$ is the output of the attention mechanism, $\boldsymbol{g}$ is the gate vector controlling information flow, $\sigma$ is the sigmoid activation function, and $E_{bg}$ is the resulting background encoding that combines temporal and attention features. This gating mechanism adaptively balances the contribution of temporal and sequence information, allowing the model to prioritize the most informative features for each variant.

Additionally, we update sequence representations with temporal information through a bidirectional information exchange:

$$\begin{aligned} S_{proj}=\text{Linear}\left( E_{seq}\left[ :,:,0,: \right] \right)\#\left( 7 \right) \end{aligned}$$

$$\begin{aligned} S_{attn},_{=}\text{MultiheadAttention}\left( S_{proj},E_{bg},S_{proj} \right)\#\left( 8 \right) \end{aligned}$$

$$\begin{aligned} S_{out}=\text{Linear}\left( S_{proj}+S_{attn} \right)\#\left( 9 \right) \end{aligned}$$

$$\begin{aligned} E_{{seq}^{bg}}=\text{Linear}\left( S_{out} \right)\#\left( 10 \right) \end{aligned}$$

where $E_{seq}$ is the sequence embedding from the sequence encoder, $E_{seq}[:,:,0,:]$ represents the first token embedding of each sequence, $S_{proj}$is the projection of this token embedding to the background encoder's hidden dimension, $E_{bg}$ is the background encoding from the previous step, $S_{attn}$ is the output of the attention mechanism where sequence projections attend to background features, $S_{out}$ is the result of a residual connection and linear transformation, and $E_{{seq}^{bg}}$ is the expanded representation (expanded along the sequence length dimension) that will be used to update sequence features with background information. This bidirectional exchange ensures that sequence features are contextualized with temporal dynamics and vice versa, creating a rich representation that captures both aspects of viral evolution.

Deep Mutational Scanning Encoder

The incorporation of experimental DMS data provides direct measurements of how mutations affect viral properties such as antibody escape, binding affinity, and expression levels. The DMS encoder integrates this experimental data with sequence representations. First, we process sequence embeddings to create a common representation space:

$$\begin{aligned} E_{combined}=\left[ \boldsymbol{E}_{\boldsymbol{target}};\boldsymbol{E}_{\boldsymbol{ref}} \right]\#\left( 11 \right) \end{aligned}$$

$$\begin{aligned} \boldsymbol{E}_{\boldsymbol{processed}}=\text{Normalize}\left( \text{Linear}\left( E_{combined} \right) \right)\boldsymbol{\#}\left( 12 \right) \end{aligned}$$

$$\begin{aligned} \boldsymbol{E}_{{target}^{proc}},\boldsymbol{E}_{{ref}^{proc}}=\text{Split}\left( \boldsymbol{E}_{\boldsymbol{processed}},\left[ 1,N_{bg} \right] \right)\boldsymbol{\#}\left( 13 \right) \end{aligned}$$

where $\boldsymbol{E}_{\boldsymbol{target}}$ *is the target sequence embedding,* $\boldsymbol{E}_{\boldsymbol{ref}}$ represents the reference sequence embeddings,$E_{combined}$is their concatenation, $\boldsymbol{E}_{\boldsymbol{processed}}$ is the processed embedding after linear projection and normalization,$\boldsymbol{E}_{{target}^{proc}}$ is the processed target embedding, and $\boldsymbol{E}_{{ref}^{proc}}$ is the processed reference embedding.

Next, we compute similarity scores between target and reference sequences to create attention weights. This attention mechanism allows the model to focus on reference sequences most similar to the target:

$$\begin{aligned} \boldsymbol{r}_{i,j,l}=\sum_{d=1}^{D_{seq}} \boldsymbol{E}_{{target}^{proc}}\left[ i,1,l,d \right]\cdot\boldsymbol{E}_{{ref}^{proc}}\left[ i,j,l,d \right]\boldsymbol{\#}\left( 14 \right) \end{aligned}$$

$$\begin{aligned} \boldsymbol{s}_{i,j,l}=\boldsymbol{r}_{i,j,l}\cdot\boldsymbol{M}_{delay}\left[ i,j,l \right]\boldsymbol{\#}\left( 15 \right) \end{aligned}$$

$$\begin{aligned} \boldsymbol{w}_{i,j,l}=\frac{\mathrm{ex}p \left( \boldsymbol{s}_{i,j,l} \right)}{\sum_{j^{'}=1}^{N_{ref}} \mathrm{ex}p\left( \boldsymbol{s}_{i,j^{'},l} \right)}\boldsymbol{\#}\left( 16 \right) \end{aligned}$$

where $\mathbf{r}_{i,j,l}$ is the dot product similarity between the target sequence and reference sequence $j$ at position $l$ for batch item $i$, $\boldsymbol{M}_{delay}[i,j,l]$ is a binary mask that prevents information leakage from future time points, and $\boldsymbol{w}_{i,j,l}$ is the attention weight derived from softmax normalization across reference sequences. For the standard DMS processing without antibody clusters or VAE, we proceed as follows:

$$\begin{aligned} \boldsymbol{X}_{dms}\in\mathbb{R}^{B\times N_{ref}\times L\times21}\boldsymbol{\#}\left( 17 \right) \end{aligned}$$

$$\begin{aligned} \boldsymbol{D}_{emb}=\text{Linear}\left( \boldsymbol{X}_{dms} \right)\in\mathbb{R}^{B\times N_{ref}\times L\times D_{dms}}\boldsymbol{\#}\left( 18 \right) \end{aligned}$$

$$\begin{aligned} \boldsymbol{D}_{trans}=\text{TransformerEncoder}\left( \boldsymbol{D}_{emb} \right)\boldsymbol{\#}\left( 19 \right) \end{aligned}$$

where $\mathbf{X}_{\mathrm{dms}}$ is the DMS data with 21 features per position (representing 20 amino acid types and deletion), $\boldsymbol{D}_{emb}$ is the embedded DMS features, and $\boldsymbol{D}_{trans}$ is the transformer-encoded DMS features.

Finally, we apply the attention weights to aggregate DMS features across reference sequences:

$$\begin{aligned} D_{weighted}=\sum_{j=1}^{N_{ref}} \boldsymbol{w}_{i,j,l}\cdot\boldsymbol{D}_{trans}\left[ i,j,l \right]\#\left( 20 \right) \end{aligned}$$

$$\begin{aligned} Y_{dms}=\text{LayerNorm}\left( D_{weighted} \right)\in\mathbb{R}^{B\times L\times D_{dms}}\#\left( 21 \right) \end{aligned}$$

where $D_{\mathrm{weighted}}$ represents the attention-weighted DMS features and $Y_{dms}$ is the final normalized DMS representation that will be integrated with other features in the model.

This approach enables the model to selectively incorporate DMS information from the most relevant reference sequences, creating a robust representation of mutation effects that informs the evolutionary prediction.

***Feature Integration and Output Layer***

We integrate evolutionary predictions with DMS features through a series of fusion layers. For models with antibody escape data:

$$\begin{aligned} \boldsymbol{F}_{dms}=\text{LayerNorm}\left( \text{Linear}\left( \left[ F_{target};\boldsymbol{Y}_{{dms}^{ab}} \right] \right) \right)\boldsymbol{\#}\left( 22 \right) \end{aligned}$$

Additional DMS features are integrated sequentially:

$$\begin{aligned} \boldsymbol{F}_{{dms}^{i+1}}=\text{LayerNorm}\left( \text{Linear}\left( \left[ \boldsymbol{F}_{{dms}^{i}};\boldsymbol{Y}_{{dms}^{i}} \right] \right) \right)\boldsymbol{\#}\left( 23 \right) \end{aligned}$$

For temporal trajectory prediction, we implement an LSTM-based output layer that generates predictions for multiple future time points:

$$\begin{aligned} \boldsymbol{X}_{{time}^{t}}=\boldsymbol{F}_{final}+\text{Embedding}\left( t \right),t\in0,1,\ldots,T-1\boldsymbol{\#}\left( 24 \right) \end{aligned}$$

The time embedding allows the model to distinguish between different prediction horizons. The LSTM processes these time-embedded features:

$$\begin{aligned} \boldsymbol{h}_{\boldsymbol{t}},\boldsymbol{c}_{t}=\text{LSTM}\left( \boldsymbol{X}_{{time}^{t}},\boldsymbol{h}_{\boldsymbol{t-1}},\boldsymbol{c}_{t-1} \right)\boldsymbol{\#}\left( 25 \right) \end{aligned}$$

$$\begin{aligned} p=\sigma\left( \text{Linear}\left( \boldsymbol{h}_{\boldsymbol{T-1}} \right) \right)\#\left( 26 \right) \end{aligned}$$

where $F_{target}$ is the output from the evolutionary prediction module for the target sequence, $\boldsymbol{Y}_{{dms}^{ab}}$ is the antibody escape DMS representation, $\boldsymbol{F}_{dms}$ is the fused representation after integrating evolutionary and antibody escape features, $\boldsymbol{Y}_{{dms}^{i}}$ is the representation of the i-th DMS feature type, $\boldsymbol{F}_{{dms}^{i}}$is the fused representation after integrating the i-th DMS feature, $\boldsymbol{F}_{final}$ is the sequence-level representation extracted from the first token position, $\boldsymbol{X}_{{time}^{t}}$ is the time-embedded representation for time point t, $\boldsymbol{h}_{\boldsymbol{t}}$ and $\boldsymbol{c}_{t}$ are the hidden and cell states of the LSTM at time step t, $p$ is the predicted prevalence probability.

This architecture enables the model to predict variant prevalence trajectories over time, capturing both immediate and longer-term evolutionary dynamics.

Training and optimization

All models were trained using the AdamW optimizer with a weight decay of 10^−2^ and an initial learning rate of 10^−4^. A linear warm-up schedule was applied for the first 300 training steps, followed by a linear decay. Mixed precision (bfloat16 or float16) was employed depending on hardware compatibility.

| Hyperparameter | Value |
| --- | --- |
| Optimizer | AdamW |
| Warm-up steps | 300 |
| Learning rate | 0.0001 |
| Weight decay | 0.01 |
| Batch size | 5 |
| Early stop patience | 10 |
| Number of axial attention layers | 3 |
| Number of transformer encoder layers | 2 |
| Number of LSTM layers | 2 |
| Number of total trainable parameters | 15.66 million |

Table S1 | Hyperparameters of DeepCoV for JN.1 era prediction

The loss function was a log-transformed, sample-weighted MSE defined as:

$$\begin{aligned} L_{reg}=\sum_{i=1}^{N} w_{i}^{\text{mask}}\cdot w_{i}^{\text{label}}\cdot\left( \log\left( \hat{y_{i}}\cdot100+1 \right)-\log\left( y_{i}\cdot100+1 \right) \right)^{2}\#\left( 27 \right) \end{aligned}$$

where $\hat{y_{i}}$ is the predicted future prevalence, $y_{i}$ is the observed ground truth at time t1 (t1= t0+30d), and $w_{i}$ is a weight derived from sequence sampling coverage masking. The mask matrix consists of binary indicators denoting whether the ground truth value $y_{i}$ is valid (i.e., total number of isolates at t1 is above a defined threshold, e.g., 100 in this study). $w_{i}^{\text{label}}$ is determined by the label $y_{i}$, to upweight the dominant variants, reducing class imbalance bias during training. To account for the heavy-tailed distribution of the target ratio and the overrepresentation of near-zero values, we applied a logarithmic transformation of the form $log\left( 100\cdot y_{i}+ 1 \right)$to the target variable. This transformation mitigates the dominance of extremely small values during training and facilitates more balanced gradient propagation throughout optimization.

Evaluation metrics

To provide interpretable and standardized performance evaluations, we computed the following metrics:

1. Mean squared error (MSE)

$$\begin{aligned} MSE=\frac{1}{n}\sum_{i=1}^{n} \left( y_{i}- \hat{y}_{i} \right)^{2}\#\left( 28 \right) \end{aligned}$$

where $y_{i}$ and $\hat{y}_{i}$ denote the true and predicted prevalence values, respectively, and $n$ is the number of data points.

2. Root mean squared error (RMSE)

$$\begin{aligned} RMSE=\sqrt{MSE}=\left. \left( \frac{1}{n}\sum_{i=1}^{n} \left( y_{i}- \hat{y}_{i} \right)^{2} \right. \right)^{1/2}\#\left( 29 \right) \end{aligned}$$

Lower RMSE values indicate closer alignment between predicted and observed variant prevalence.

3. Pearson correlation coefficient *r*

$$\begin{aligned} r=\frac{\sum_{i=1}^{n} \left( y_{i}-\bar{y} \right)\left( \hat{y_{i}}-\bar{\hat{y}} \right)}{\sqrt{\sum_{i=1}^{n} \left( y_{i}-\bar{y} \right)^{2}}\cdot\sqrt{\sum_{i=1}^{n} \left( \hat{y_{i}}-\bar{\hat{y}} \right)^{2}}}\#\left( 30 \right) \end{aligned}$$

where $\bar{y}$ and $\bar{\hat{y}}$ represent the means of the true and predicted values, respectively.

Generalization

Updated JN.1 era prediction

For the updated JN.1 prediction, the main model—originally trained on data collected before 1 October 2023—was retained, while the test dataset was expanded to include sequences submitted up to 16 May 2025. Unique RBDs were renamed according to their mutations relative to reference lineages: WT, Alpha, Beta, Delta, Gamma, Eta, BA.1, BA.2, BA.5, BF.7, BQ.1.1, XBB, XBB.1.5, EG.5, HK.3, Flip, BA.2.86, JN.1, KP.2, KP.3, XEC, LF.7, LP.8, NB.1, NB.1.8.1, XFG, and XFH. The major-strain-specific test set comprised RBDs from JN.1, KP.2, KP.3, LF.7, LP.8, and NB.1.8.1.

Spike model

For each unique spike sequences, cluster index and cluster name are assigned for further usage. Unique spike were renamed according to their relative mutations relative to the their parental lineage references: WT, Alpha, Beta, Delta, Gamma, Eta, BA.1, BA.2, BA.5, BF.7, BQ.1.1, XBB, XBB.1.5, EG.5, HK.3, “FLip” (XBB+S486P+L455F+F456L), BA.2.86, JN.1, KP.2 and KP.3. For the sequencing embedding, considering the input restrictions of the ESM-MSA-1b, only the first 1,023 amino acids of the spike protein were truncated. Considering the relatively minor contribution of the tail of the C-terminal region in the evolution of SARS-CoV-2, it is an acceptable compromise. The other data processing and model architecture are consistent with those of the main model in all other aspects.

| Hyperparameter | Value |
| --- | --- |
| Optimizer | AdamW |
| Warm-up steps | 300 |
| Learning rate | 0.0001 |
| Weight decay | 0.01 |
| Batch size | 5 |
| Early stop patience | 3 |
| Number of axial attention layers | 3 |
| Number of transformer encoder layers | 2 |
| Number of LSTM layers | 2 |

Table S2 | Hyperparameters of Spike model

XBB era model

Data prior to September 1, 2022, were used for the training and validation sets, while data collected afterward were reserved for the test set. To accommodate the reduced training data while maintaining model performance, we decreased the number of self-supervised transformer layers in the MSA-proportion fusion module from three to two. The other data processing and model architecture are consistent with those of the main model.

| Hyperparameter | Value |
| --- | --- |
| Optimizer | AdamW |
| Warm-up steps | 300 |
| Learning rate | 0.0001 |
| Weight decay | 0.01 |
| Batch size | 5 |
| Early stop patience | 10 |
| Number of axial attention layers | 2 |
| Number of transformer encoder layers | 2 |
| Number of LSTM layers | 2 |

Table S3 | Hyperparameters of XBB model

Continuous model

For a given t₀, the model predicts variant growth over the next 0-60 days. In terms of model architecture, the output layer was modified from two fully connected layers to an LSTM layer. For the loss function, we introduced an additional per-day weighting term $w_{k}^{\text{day}}$ into the regression loss:

$$\begin{aligned} \mathcal{L}_{\text{cn}}=\sum_{i=1}^{N} \sum_{k=1}^{60} \left[ w_{i,k}^{\text{mask}}\cdot w_{i,k}^{\text{label}}\cdot w_{k}^{\text{day}}\cdot\left( \log\left( \hat{y_{i,k}}\cdot100+1 \right)-\log\left( y_{i,k}\cdot100+1 \right) \right)^{2} \right]\#\left( 31 \right) \end{aligned}$$

$w_{i,k}^{\text{mask}}$ filters out invalid timepoints and $w_{i}^{\text{label}}$ is associated with the label $y_{i,k}$, to reduce class imbalance bias. $w_{k}^{\text{day}}$ is a temporal weight associated with prediction k, shared across all samples. It is designed to emphasize time points within the 60-day prediction window that are both more informative and label-stable. To implement this, we constructed a Gaussian weighting vector over the time axis, centered at μ=30 with a standard deviation σ=10, assigning greater importance to mid-range days. The other data processing and model architecture are consistent with those of the main model.

| Hyperparameter | Value |
| --- | --- |
| Optimizer | AdamW |
| Warm-up steps | 300 |
| Learning rate | 0.0001 |
| Weight decay | 0.01 |
| Batch size | 5 |
| Early stop patience | 10 |
| Number of axial attention layers | 3 |
| Number of transformer encoder layers | 2 |
| Number of LSTM layers (proportion embedder) | 2 |
| Number of LSTM layers (output) | 1 |

Table S4 | Hyperparameters of continuous model

Benchmarking approaches

Growth advantage estimation

The algorithm for calculating growth advantage was adapted from Chen et al., and the daily sequence data were sourced from the GISAID database^11^. Specifically, a logistic regression model was employed to fit the daily frequency of samples for the concerning RBD cluster to estimate the logistic growth rate *a* and the midpoint *t0* of the sigmoid curve. The growth advantage was defined as e^a × g^ − 1, where g, the generation time, equals to 7 days, and *a* represented the growth rate derived from the logistic model fitting. Confidence intervals were computed with α=0.95.

EVEscape benchmarking

The algorithm for calculating growth advantage was adapted from Thadani et al. ^12^. We first computed the mutations of all RBD sequences in the full test set relative to the wild-type reference. Following the approach described in the original EVEscape publication, we aggregated EVEscape scores based on these relative mutations to obtain a composite score for each RBD sequence.

E2VD benchmarking

The E2VD models were retrained using the publicly available ESM-2 model as the pre-trained backbone^13^. Candidate unique RBD sequences served as input to the model, with outputs generated on a five-fold cross-validated test set. We benchmarked the E2VD framework by aggregating predictions from its three submodules: ACE2 binding affinity, viral expression efficiency, and antibody escape potential. Model outputs were generated on a five-fold cross-validated test set (JN.1-era RBD variants) and averaged across folds for the binding and expression tasks. Escape scores were computed from the last prediction run of the optimization cycle. Following the original study, the model was tasked with predicting escape scores against the BD57-0129 antibody for immune escape evaluation. To prioritize variants with potential fitness and immune evasion advantages, we applied an asymmetric scoring scheme informed by prior biological knowledge: deviations in expression and ACE2 binding were penalized only when falling below functional thresholds, whereas antibody escape was positively weighted when exceeding a permissive cutoff. The final E2VD score was computed as the sum of exponentiated deviations from these empirically defined thresholds:

$$\begin{aligned} \text{E2VD}=e^{\min\left( \text{expr}-0.7, 0 \right)}+e^{\min\left( \text{bind}-0.25, 0 \right)}+e^{\max\left( \text{escape}-0.5, 0 \right)}\#\left( 32 \right) \end{aligned}$$

***Fitting model of DMS***

We developed a statistical predictive framework leveraging deep mutational scanning (DMS) and antibody neutralization data. This model assumes that the selective advantage of a new variant *v* in a given population is directly related to its capacity to escape neutralization. The escape potential is determined by two factors: (1) the composition of antibodies in the host immune landscape, and (2) the predicted neutralization efficacy of each antibody against *v*.

$$\begin{aligned} I\left( A,\nu\right)=\sum_{a\in A} \frac{1}{n\left( a,\nu\right)}\#\left( 33 \right) \end{aligned}$$

Here, $I\left( A,\nu\right)$ represents the cumulative neutralization inverse scores across antibody set A, and $n\left( a,\nu\right)$ denotes the neutralization titer (e.g., IC₅₀) of antibody a against variant ν. Because experimental neutralization data is unavailable for all possible variant–antibody combinations, we estimate neutralization changes using DMS-derived escape scores.

$$\begin{aligned} n\left( a,\nu\right)=g\left( S_{a}\left( \mathbf{m}\left( \nu,R \right) \right) \right)n\left( a,R \right)\#\left( 34 \right) \end{aligned}$$

where R is a reference strain (e.g., D614G, BA.2, BA.5), and $\mathbf{m}\left( \nu,R \right)$ denotes the set of RBD amino acid mutations that differentiate variant ν from R. The function $S_{a}\left( \mathbf{m} \right)=\left\{ S_{a}\left( m_{i} \right) \right\}$ denotes the per-site DMS escape scores for antibody a, and g(·) is a transformation function applied to escape scores. To ensure interpretability and monotonicity, we require g(x) ≥ 1 (as higher DMS escape implies weaker neutralization) and set g(0) = 1, with g(1) → ∞.

We then define the relative immune escape advantage of variant ν compared to reference R as:

$$\begin{aligned} Q_{e}\left( \nu;A,R \right)=-\log\left( \frac{I\left( A,\nu\right)}{I\left( A,R \right)} \right)=\log I\left( A,R \right)-\log I\left( A,\nu\right)\#\left( 35 \right) \end{aligned}$$

This yields a dimensionless immune escape score relative to a reference strain, quantifying the extent to which a variant evades population-level immunity. For the transformation g(x), we adopt the following parametric form to satisfy the above properties:

$$\begin{aligned} g\left( x \right)=\exp\left[ k_{M}\cdot\max\left( x \right)+k_{s}\cdot\sum\left( x \right) \right]\#\left( 36 \right) \end{aligned}$$

where $k_{M}$ and $k_{s}$ are tunable parameters. This formulation ensures that escape effects are additive and non-negative, while amplifying both hotspot and distributed escape contributions. To train the model, we compiled five curated neutralization datasets covering reference-to-variant transitions such as BA.2 to BA.5 and D614G to BA.1, encompassing a range of mutation loads and escape profiles. Fitting a linear model to these datasets yielded robust and interpretable parameters.

Benchmarking settings

Timelines of major variant detection by different predictive models

To assess the timeliness of variant prioritization across different computational frameworks, we tracked the earliest time points when major RBD variants (JN.1, KP.2, and KP.3) were flagged as high-risk by multiple scoring models, including DeepCoV, EVEscape, and E2VD. For each method, we determined the earliest date a given strain entered the top-N ranking (e.g., top 10 or top 3) based on model-specific scores. Specifically, DeepCoV rankings were derived from the model-predicted target proportion using data available at date t₀. For EVEscape and E2VD, strain scores were ranked within a ±1-month temporal window centered at each submission date to mimic realistic evaluation scenarios.

Benchmarking of Top-k Variant Ranking Predictions

To enable a fair comparison of different predictive methods for identifying dominant SARS-CoV-2 RBD variants during the JN.1 era, we implemented a dynamic retrospective top-*k* ranking comparison against several baseline methods. The evaluation was conducted on temporally stratified test data spanning October 2023 to September 2024. We first defined ground-truth dominant variants (*top-n_truth*) based on their observed proportions at prediction horizon *t₁*, using a 30-day window of metadata (submit_date ∈ [t₀−30, t₀]) to restrict candidate RBDs to those actively circulating at the time of prediction. For each time point *t₀*, candidate variants were ranked by predicted prevalence using DeepCoV, by growth advantage, and by baseline functional scores from EVEscape and E2VD. We evaluated using two criteria:

(1) ***Prediction success rate*** (for cases where the number of true dominant variants = 1): defined as the proportion of time points where the top-ranked predicted variant exactly matched the observed dominant variant at *t₁*, based on its prevalence.
(2) ***Mean Jaccard Index*** (for cases where the number of true dominant variants > 1): the Jaccard Index is defined as

$$\begin{aligned} Jaccard\left( A,B \right)=\frac{\left| A\cap B \right|}{\left| A\cup B \right|}\#\left( 37 \right) \end{aligned}$$

where *A* and *B* denote the sets of top-*n* predicted and observed variants, respectively. The Mean Jaccard Index was calculated as the temporal average of the Jaccard Index across all evaluation points, quantifying the overlap between predicted and true dominant sets. The top-*k* predictions were computed for a range of values k. Evaluation was conducted independently for each method across all time points.

Aggregate TopK comparison

We assessed model performance using rank-based evaluations, acknowledging that the output scores from different models vary in scale and interpretation. For each method, RBD variants were ranked in descending order according to their predicted scores, and the top *k* variants were compared against a predefined set of *n* dominant variants, those with the highest observed prevalence. Evaluation metrics, including recall (true positive rate) and FDR, were computed at each value of *k*. These metrics collectively provided a comprehensive evaluation of each method’s ability to prioritize high-fitness, high-impact variants. When evaluating the model trained on data prior to 1 October 2023 (JN.1 era) and its ablated variants, we used HK.3, BA.2.86, JN.1, KP.2, and KP.3-related variants as the evaluation sequence set.

Dynamic assessment under variable prevalence threshold

Compared to other methods, our model produces prediction scores with intrinsic quantitative interpretations, enabling direct linkage to real-world variant prevalence. In practice, the identification of dominant strains depends on the definition of dominance, which may vary by context or policy need. This means our framework can flexibly infer potential dominant variants under different epidemiological expectations.

To systematically evaluate the sensitivity of our model across varying definitions of variant dominance, we performed a dynamic benchmarking analysis using a three-stage framework (T1–T3). Specifically, we defined the following key timepoints for each variant based on surveillance data:

- **T1:** the earliest timepoint at which the cumulative number of observed sequences for a variant exceeds 10, indicating initial emergence.
- **T2:** the first date when the variant exhibits consistent growth advantage (>30%) over a 7-day sliding window, provided it had accumulated at least 100 sequences.
- **T3:** the earliest date at which the variant’s relative proportion among all sequences surpasses a predefined threshold.

For each variant and prevalence threshold, we determined whether the model successfully predicted that the variant’s future proportion would exceed the given T3 threshold. Based on these, we computed recall and FDR for each threshold. For XBB era prediction, the candidates were constrained to XBB, EG.5, BQ.1 and HK.3 subvariants.

Growth trajectory reconstruction and Prevalence regional heterogeneity analysis

Growth trajectory reconstruction

To evaluate the temporal dynamics and predictive accuracy of our model across key SARS-CoV-2 lineages, we visualized the predicted and observed relative abundances of major RBD or spike variants over time. For each lineage, model predictions of future proportions (output of target ratio at t_1_) were aligned to their corresponding start dates (t_0_) and binned biweekly. Observed contemporaneous proportions (target ratio at t_0_) were used for reference.

Line plots were generated to display predicted trajectories (dashed lines), observed proportions (solid lines), and prediction uncertainty (±1 SD as shaded ribbons) across locations. Variants were labeled using standardized nomenclature derived from mutational mappings. We computed temporal prediction error metrics, including RMSE and MAE, to quantify model fidelity across time points. These metrics were evaluated both on major variants and the full test set.

Prevalence regional heterogeneity analysis

For several well-known variants exhibiting regional differences in prevalence, we separately modeled their growth trajectories using representative data from individual continents and visualized their predicted dynamics. In addition, to assess the spatiotemporal dynamics of variant emergence, we evaluated the relative ranking of four representative RBD variants across major geographic regions. The test dataset consisted of model outputs spanning from 1 September 2022. For each variant, predictions were grouped by date (*t₀*) and location, and variants were ranked according to their predicted prevalence at the forecast horizon (*t₁*).

In silico mutational hotspots scanning

Pseudo single-residue mutational dataset generation

We constructed comprehensive libraries of single amino acid substitutions on representative reference strains. Aligned regions corresponding to the RBD (residues 331–531) or NTD (positions 14–305) were extracted from multiple sequence alignments of spike protein sequences. For each position in the aligned region, and for each reference strain (e.g., XBB.1.5, JN.1, KP.2), all 19 possible single amino acid substitutions were introduced. In the region of NTD, extra deletions are also introduced for each position. To simulate synthetic surveillance records, we selected a defined time window (e.g., May 1 to June 1, 2023, for XBB.1.5) and globally assigned each mutant to every retained their empirical prevalence values; unobserved mutants were assigned a prevalence of zero. The combined dataset was used for downstream mutational hotspot scanning and predictive evaluation.

In silico mutational scanning and visualization of variant-specific fitness hotspots

In silico single-mutant scanning of the SARS-CoV-2 RBD or NTD region were conducted on various emerging variant backbones (e.g., XBB.1.5, KP.3, LF.7). For reference strain JN.1，KP.3 and LF.7, all single amino acid substitutions were evaluated using trained JN.1-era predictive model, yielding a fitness score for each synthetic mutant. For reference strain XBB, XBB.1.5 and EG.5, residue substitutions were evaluated using trained XBB-era predictive model. For each mutation, a normalized contribution score was calculated by subtracting the amino acid–specific global average score over time to control for residue bias. Only mutations with a positive differential score were retained as candidates for fitness advantage. To summarize mutation-level signals into position-level insights, we computed average contribution scores across all substitutions at each site and visualized them using a smoothed line plot with positional annotations. Additionally, we generated amino acid logo plots on the top N (e.g., 10) highest-scoring positions, where the height of each letter corresponded to the magnitude of the fitness contribution for that substitution.

Ablation studies

To assess the contribution of different components to the model's predictive performance, we conducted systematic ablation studies by removing specific functional modules:

- **Sequence encoder ablation (ESM-2)**: To evaluate the effect of the MSA-based encoder, we replaced the ESM-MSA-1b with the ESM-2 (150M) model for sequence embedding. All other architectural components were kept unchanged.
- **No-DMS model**: All DMS-derived features were removed, and the model was trained using only the amino acid sequences and associated epidemiology data. Two feed-forward layers are used in place of the DMS module to achieve the same dimensional transformation.
- **No background model**: Only the target strain and its associated 180-day historical prevalence data were used. No background sequences were included. Sequence features were encoded using the ESM-2 model, and historical prevalence signals were integrated via an LSTM, followed by a Transformer-based feature aggregator.
